# BioMeme: biological response memory for virtual cells

**DOI:** 10.64898/2026.11.07.765043

**Authors:** Jia Bi, Chenyang Zhu, Haochen Liu, Yufei Wang, Ting Liu, Xinqi Bao, Runshan Hu, Jeyan Thiyagalingam, Steve R. Gunn

**Affiliations:** Scientific Computing Department, Science and Technology Facilities Council, Rutherford Appleton Laboratory, Didcot, OX11 0QX, UK; School of Electronics and Computer Science, University of Southampton, Southampton, SO17 1BJ, UK; School of Electronics and Information, Northwestern Polytechnical University, Xi’an, 710072, China; Division of Information Science and Engineering, KTH Royal Institute of Technology, Stockholm, SE-100 44, Sweden; Jimei University, Jimei District, Xiamen, Fujian 361021, China

**Author notes:** Contributing authors.

## Abstract

Virtual cells predict responses to unmeasured perturbations, but neural models absorb measurements into weights, require retraining and cannot cite supporting experiments. Here we introduce biological response memory, replacing a learned state-to-response layer with a memory of measured experiments. BioMeme predicts next-dose responses in a new cell type from measured lower-dose states, recalling similarly responding compounds in that cell and the same compound in a reference cell line. Each prediction is an exact weighted sum of named measurements, and memory updates with one linear solve. In all nine preregistered comparisons across three screens, memory predicted response direction better than the learners it replaced. In U2OS, BioMeme outperformed input-matched ridge, adapted chemCPA and a neural layer in memory’s place, and matched a full-panel network reading reference responses directly (error within 3.4%). Transfer was compound-specific, lowering error by 21.2% against at most 1.8% for wrong compounds, and replicated in reserved MCF7 data. With 20 references, fitting took seconds on one CPU thread, at least 420 times faster than neural tuning and fitting. Adding its prediction to one measured well cut that well’s error by 27–31%. Biological response memory establishes a design principle for virtual cells any laboratory can update, inspect and run.

## Introduction

Knowing how a cell responds to a compound, at which dose and in which cell type, underpins drug discovery and cell biology. Testing 1,000 compounds at three doses in 10 cell lines requires 30,000 conditions before replication. Most combinations remain unmeasured, and every new cell type widens the gap.

Virtual cells aim to close this gap by predicting untested perturbations from biological measurements^[1–3]^. Atlases span millions of transcriptional, morphological and protein profiles^[4–9]^. Multi-cell-line Cell Painting and pooled single-cell screens measure how compound responses vary across cell types^[10,11]^. Useful predictions connect these atlases with local experiments^[2]^. Screening laboratories need to choose wells, toxicologists concentrations, and chemical biologists compounds with shared mechanisms.

Predictions degrade outside training conditions, and simple mean or linear predictors often match deep-learning models ^[12–15]^. Leading approaches in the 2025 Virtual Cell Challenge combined neural networks with conventional statistics^[16,17]^. Generative models approach replicate agreement for new combinations of conditions represented in a training atlas^[18]^. Latent-variable, optimal-transport and graph-based models learn mappings between states and perturbations^[19–25]^, while image-based and multimodal models extend prediction to morphology ^[26–30]^. The 3-billion-parameter Tahoe-x1 foundation model was pretrained across 128 GPUs^[31]^. Learning compresses experiments into parameters and leaves individual contributions implicit. Dose screens also supply direct evidence: the query compound at a lower dose, other compounds in the target cell and the same compound in a reference cell. Predicting the next dose from this evidence is the virtual-cell question in its everyday screening form: a measured cell state and a perturbation, mapped to the resulting state^[1]^. Laboratories where robots and AI agents plan experiments^[32,33]^ need predictors that absorb such measurements, expose their experimental support and fit locally.

Several methods consult observations at prediction time. LangPert and retrieval guided by large language models select and average measured perturbations, while MechPert adds regulatory agreement^[34–36]^. scPRAM combines latent perturbation vectors from matched untreated cells, and PT-RAG retrieves gene embeddings from the control-cell state^[37,38]^. Stack conditions on measured cell populations, while response memories and cross-context kernel regression combine observations with learned predictions^[14,39,40]^. Rhaister learns drug–panel relationships across reference contexts and combines panel measurements in a new context using ridge weights^[41]^. BioPert supplies a compound’s reference-cell measurement and the target’s untreated profile to a neural network ^[42]^. Tensor completion and deep learning also borrow across drug and cell axes^[43,44]^. Dose escalation offers another cue: how the query compound has already acted in the target cell. We use this treated state to connect target-cell memory with same-compound transfer while preserving the contributing experiments’ identities.

Here we introduce biological response memory, replacing a learned state-to-response layer with a memory of measured experiments, and implement it as BioMeme (Fig. 1). Dose responses depend on both the compound and the cell, motivating two routes. Kernel memory recalls other compounds’ residual responses in the target cell through a Gaussian-process posterior mean, using the query’s lower-dose state to judge relevance. Reference transfer supplies the same compound’s reference-cell dose response through a strength gate and ridge correction. A regularized solve turns state similarities into signed experiment weights. Expanding calibration, memory and transfer gives a weighted sum of named measurements for the complete prediction. Memory shares inputs and outputs with the learners it replaces. New responses enter through a linear solve.

**Fig. 1.**
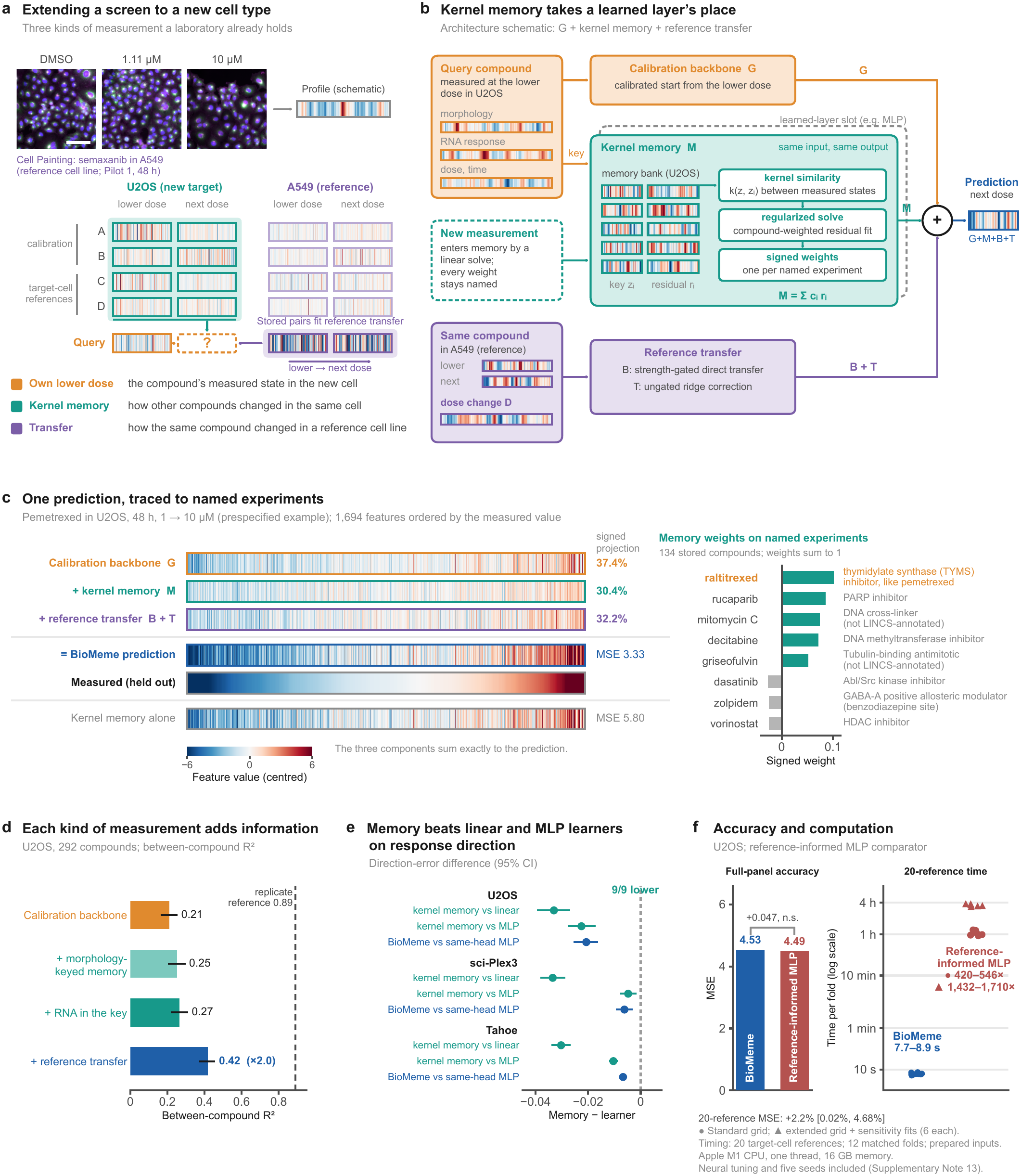
| BioMeme recalls measured biological responses. **a,** Extending a screen to a new cell type. Semaxanib images show A549 Pilot 1 at 48 h (DNA, blue; endoplasmic reticulum (ER), green; actin/Golgi/plasma membrane (AGP), magenta; scale bar, 100 µm; display ranges, Supplementary Table 28). Images illustrate the design; analysed profiles are Batch 2. The image-to-profile symbol is schematic. **b,** The calibration backbone *G* is augmented by kernel memory *M* and reference transfer. Compound-weighted fitting gives signed residual weights. *B* denotes strength-gated direct transfer; ridge correction *T* is ungated. *z_i_*denotes an experimental key, *r_i_*its stored residual and *c_i_*its signed weight (*cz* in Eq. 4). A linear solve updates memory. **c,** Prespecified pemetrexed example: U2OS, 48 h, nominal 1 *→* 10 µM. All 1,694 features share measured-value order and colour limits *±*6. Percentages are signed projections onto the centred prediction, subtracting the calibration centre from *G*; components sum before display clipping. Bars show five largest positive and three most negative weights among 134 compounds; all weights sum to one. Negative weights arise from regularization, without biological interpretation. **d,** Between-compound *R*^2^, after removing each scoring group’s mean prediction error, for 292 U2OS compounds in twelve groups. The replicate-derived reference assumes independent measurement noise. **e,** Direction-error differences: kernel memory versus linear or multilayer perceptron (MLP) learners without transfer; BioMeme versus the same-head MLP (memory replaced by an MLP with the same transfer head). All nine passed two-sided centred-bootstrap tests with Holm correction within screen families. **f,** Full-panel MSE (left); twenty-reference workflow times from prepared inputs in twelve matched folds (right). Same CPU, one numerical thread per process; MLP workers ran concurrently, BioMeme serially. Neural times include tuning, five seeds and, in six folds, extensions and original-configuration sensitivity fits (Supplementary Note 13). At twenty references, BioMeme’s MSE was 2.20% higher than the reference-informed MLP’s (95% interval, 0.02–4.68%); direction error did not differ significantly (Supplementary Table 52). Intervals are 95% whole-compound bootstrap intervals (10,000 draws). Source data are provided with this paper.

We evaluated BioMeme on three public screens spanning morphology and single-cell transcriptomes, with predictions locked before scoring and statistics recomputed by separately written code. Memory predicted direction better than the learners it replaced in all nine preregistered comparisons. In U2OS, BioMeme doubled the calibration backbone’s explained variation and outperformed a mean-response baseline, input-matched ridge, adapted chemCPA^[21]^ and an MLP in memory’s place. With the full U2OS panel, BioMeme matched a tuned network that reads reference responses directly, and reserved MCF7 data replicated this parity. In 241 U2OS compounds, matched reference compounds lowered mean squared error (MSE) by 21.2%, compared with 0.7% and 1.8% for wrong compounds matched for response energy or cell-count trajectory. Full transfer replicated in reserved MCF7 data with 25% lower MSE. Every prediction expanded exactly into weighted named measurements. Deleting the five most heavily weighted compounds changed predictions 4.5 times as much as random deletion, and experiments with the five largest positive weights were 3.3-fold enriched for shared mechanisms or targets. Fitting took seconds on one CPU thread, compared with 1–4 h for neural tuning and fitting. Adding BioMeme’s prediction to one measured well lowered single-well error by 27–31%.

For screening laboratories, useful references profile the same compound, batch and doses. Developers can replace a learned response layer with memory, update it in closed form and trace its contributions. Prepared tasks on three screens support benchmarking, and named compounds give chemical biologists hypotheses about shared mechanisms. Primary or patient-derived cells and Cell Painting toxicology offer applications^[45]^. A CPU implementation and Library of Integrated Network-based Cellular Signatures (LINCS) tutorial accompany this paper. Biological response memory makes measured experiments available for virtual cells to recall, inspect and update, from local screens to automated laboratories.

## Results

### BioMeme predicts responses at an untested dose from three kinds of measurement

BioMeme predicts next-dose responses from related target-cell states and the same compound’s reference-cell response (Fig. 1a). The query’s lower-dose profile anchors prediction and conditions recall. Signed weights identify contributing experiments (Fig. 1b,c). Reference transfer adds strength-gated direct transfer (*B*) and a ridge correction (*T*); removing both leaves kernel memory.

Query targets were withheld, and the reference cell line was measured at both doses. Separate compounds fitted calibration or supplied target-cell references. Preregistered evaluations covered U2OS Cell Painting, sci-Plex3 transcriptomes (MCF7 and K562 targets) and Tahoe-100M (Tahoe; 10 target lines from 10 tissues) ^[5,6,46]^.

Each route reduced MSE when added to the other. Linear and one-hidden-layer multilayer perceptron (MLP) learners replaced memory with identical inputs and outputs. The same-head MLP retained BioMeme’s transfer head. Memory lowered direction error, one minus the cosine similarity between centred predicted and measured profiles, in all nine preregistered comparisons (Fig. 1e).

Adding these sources increased explained variation across 292 U2OS compounds (1,168 predictions). Between-compound *R*^2^, which removes each scoring group’s mean prediction error, rose from 0.21 for the backbone to 0.25 with morphology-keyed memory and 0.27 after adding RNA. Reference-cell responses raised it to 0.42, against a noise-based replicate reference of 0.89 (Fig. 1d). Calibration plus transfer alone reached 0.401 (0.354–0.440; Supplementary Table 70). BioMeme removed 35.5% (31.6–39.4%) of the backbone’s reducible error (error above replicate noise); MCF7 replication gave 38.2% (33.3–43.1%; Supplementary Fig. 7 and Supplementary Table 21).

### Same-compound dose responses recur across cell types

Morphological dose-response correlations between cell types retained 49% of the same-cell between-batch correlation. A549 images and profiles show changes recurring in U2OS (Fig. 2a,b; Supplementary Fig. 4). Across 80 compounds, A549 Pilot 1 dose responses correlated with U2OS Batch 2 at 0.189 and A549 Batch 2 at 0.386. Both comparisons spanned batches. Their ratio was 0.489 (95% confidence interval, 0.336–0.620), exceeding the preregistered threshold of 0.30 (Holm-adjusted *P* = 0.0095; Fig. 2c,e; Table 1). The ratio was 0.498 after excluding shape, density and location features and 0.360 after removing each set’s first principal component, showing recurrence beyond these feature groups and the dominant response axis. Removing features with zero change in either profile left the ratio at 0.493 (0.339–0.624; Supplementary Table 11). Requiring all target-dose wells at both steps to reach half their plate’s solvent-only dimethyl sulfoxide (DMSO) median cell count left 30 of 80 compounds. Their ratio was 0.269 (0.043–0.436), above zero but below the 0.30 threshold preregistered for all 80 (Supplementary Note 10.8). Within Batch 2, matched A549–U2OS compound pairs were more concordant than mismatched pairs (median correlation, 0.31 versus 0.04; Fig. 2f) and than wrong compounds with the closest A549 cell-count trajectory (0.31 versus 0.09; mean paired difference 0.21, 0.14–0.27; Supplementary Table 68). Adjusting dose-response profiles for cell-count changes retained a cross-cell/between-batch ratio of 0.482 (0.289–0.664; Supplementary Table 68).

**Fig. 2.**
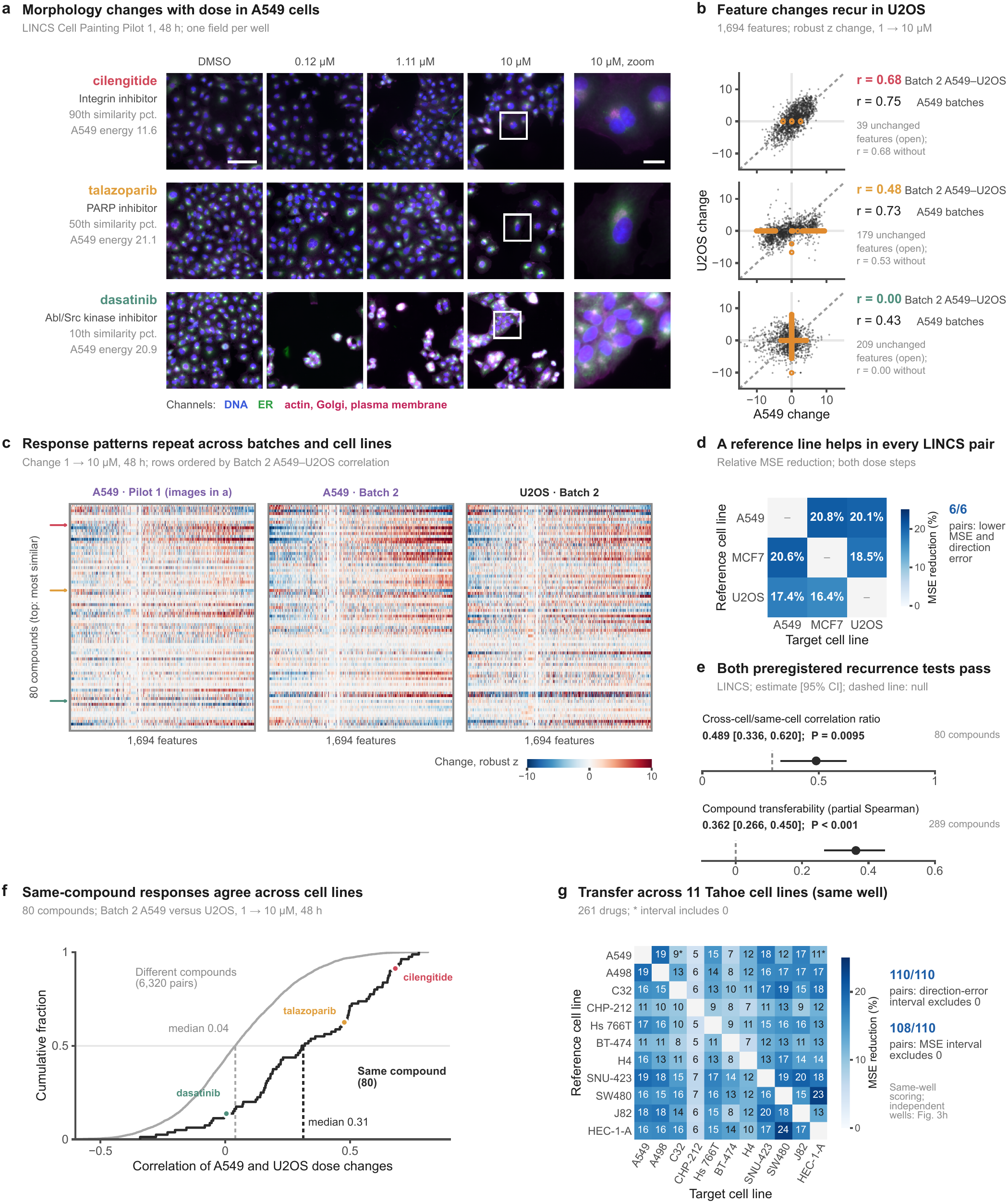
| Dose-response patterns recur across cell types. **a,** A549 Pilot 1 images at 48 h for prespecified compounds at the 90th, 50th and 10th percentiles of Batch 2 A549–U2OS similarity among strong A549 responders. Tiles show field 1 centres; zooms centre on the nearest nucleus. DNA, blue; ER, green; AGP, magenta. Pilot 1 used 0.12 and 1.11 µM for nominal 0.1 and 1 µM doses. Display ranges are shared. Scale bars, 100 µm (fields) and 20 µm (zooms). **b,** Batch 2 A549 versus U2OS feature changes, nominal 1 *→* 10 µM at 48 h; “A549 batches” compares Pilot 1 and Batch 2. Points are 1,694 features; axes span *−*15 to 15. Open orange circles mark zero-change features; correlations labelled “without” exclude these features (Supplementary Data 44). **c,** Dose changes for 80 compounds in Pilot 1 A549, Batch 2 A549 and Batch 2 U2OS. Rows descend by Batch 2 A549–U2OS correlation; features follow the first A549 principal-component loading. Display limits, *±*10 robust *z*; arrows identify compounds in **a**. **d,** Relative MSE reduction from reference transfer in six directed LINCS cell-line pairs, pooled over dose steps and times. This prespecified exploratory comparison uses nominal 95% intervals; MCF7 targets are development data. **e,** Registered recurrence tests: cross-cell/same-cell correlation ratio across both dose steps (80 compounds; threshold, 0.30) and partial Spearman association of compound transferability (289 compounds). Holm-adjusted *P* values use one-sided whole-compound bootstrap testing for the ratio and stratified permutation testing for transferability (10,000 draws each). Dashed lines show null thresholds. **f,** Batch 2 A549–U2OS dose-change correlations for 80 matched and 6,320 mismatched pairs, reusing the same compounds. **g,** Same-well Tahoe transfer across 11 lines and 261 drugs shared by all lines. Asterisks mark MSE intervals including zero; counts denote intervals excluding zero. Heatmap percentages are rounded. Fig. 3h evaluates independent wells with A549 as reference. Intervals are 95% whole-compound bootstrap intervals. Source data are provided with this paper.

**Table 1.**
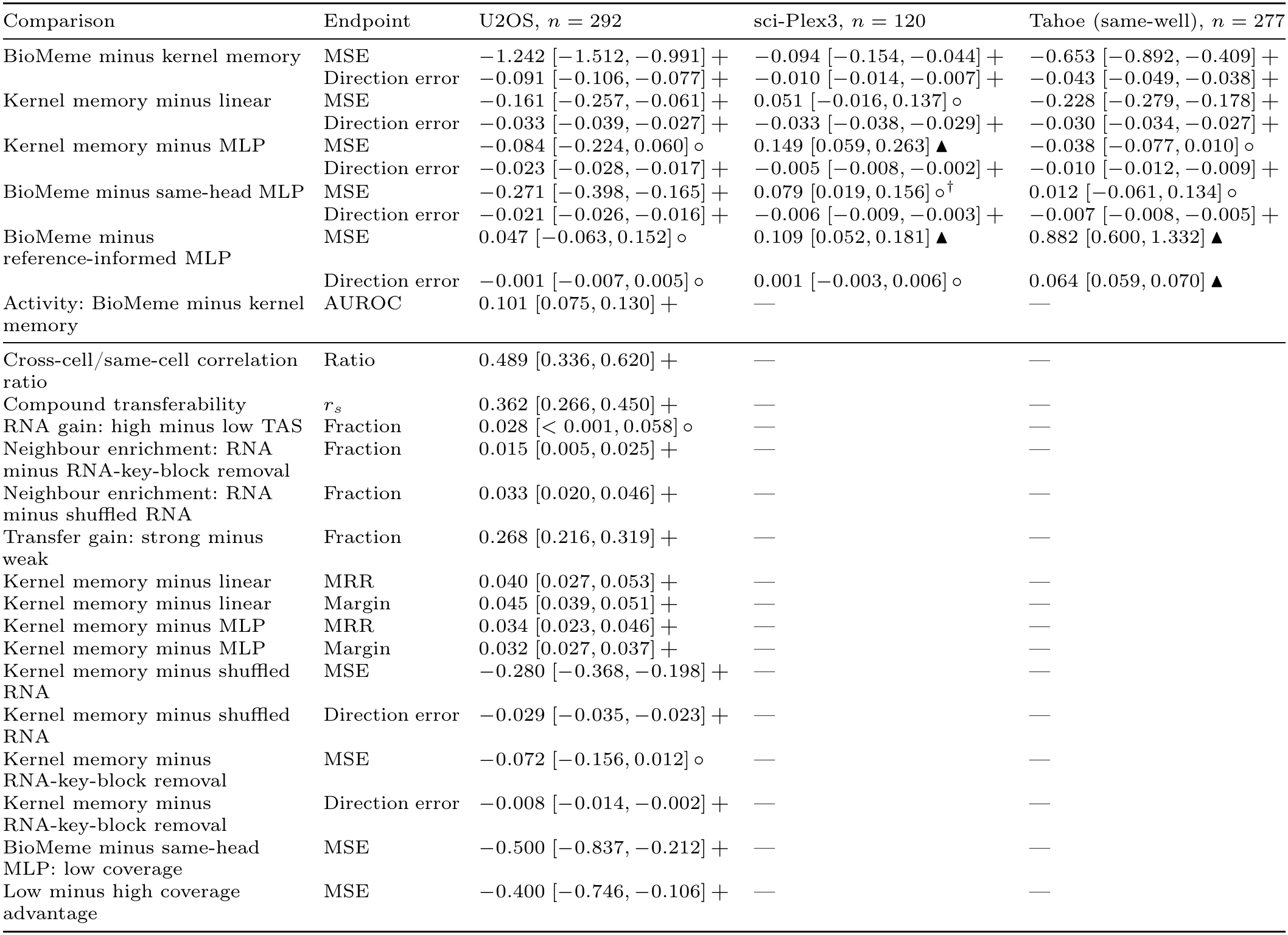
| Preregistered tests in three screens. **notes.** Entries are estimates and paired whole-compound 95% percentile intervals (10,000 draws). +: favourable; *◦*: not significant; ▴: comparator favoured. Negative loss differences and positive AUROC, identification or enrichment differences favour the first model. Symbols follow within-family Holm decisions, except the activity test and two coverage rows, which follow their registered one-sided rules. The correlation ratio and compound transferability form one Holm family; the secondary biological hypotheses form another. Both use one-sided tests at *α* = 0.025. The correlation ratio uses 80 compounds and a threshold of 0.30; compound transferability uses 289; neighbour enrichment uses 224 annotated compounds. RNA-key-block removal retains calibration geometry fitted with RNA. The high-minus-low-TAS RNA-gain interval is unadjusted; the test was not significant (unadjusted and Holm-adjusted *P* = 0.0251; *α* = 0.025; Supplementary Table 11). The sequential budget and five-reference substitution tests, including stopping decisions, are in Supplementary Table 7. Tuned-kernel supplements for U2OS and sci-Plex3 are in Supplementary Tables 7 and 8. Tahoe rows involving reference information use same-well targets; Fig. 3h and Supplementary Table 29 score the same predictions against independent replicate wells. Tahoe all-reference comparisons use the tuned model; **Tahoe MSE differences and intervals are multiplied by** 10^3^. Other MSE values retain native units. A dash denotes a test not specified for that screen. AUROC, area under the receiver operating characteristic curve; TAS, transcriptional activity score; MRR, mean reciprocal rank; margin, cosine-similarity identification margin. Test families are defined in Supplementary Note 5; MCF7 replication and supplementary analyses are in the Supplementary Information. *†*: the nominal interval favours the comparator but the Holm decision is not significant. With seed-averaged neural predictions, this comparison favours the MLP on MSE (Holm *P* = 0.047; Supplementary Table 32).

| Comparison | Endpoint | U2OS, *n* = 292 | sci-Plex3, *n* = 120 | Tahoe (same-well), *n* = 277 |
|---|---|---|---|---|
| BioMeme minus kernel memory | MSE | −1.242 [−1.512, −0.991] + | −0.094 [−0.154, −0.044] + | −0.653 [−0.892, −0.409] + |
|  | Direction error | −0.091 [−0.106, −0.077] + | −0.010 [−0.014, −0.007] + | −0.043 [−0.049, −0.038] + |
| Kernel memory minus linear | MSE | −0.161 [−0.257, −0.061] + | 0.051 [−0.016, 0.137] ∘ | −0.228 [−0.279, −0.178] + |
|  | Direction error | −0.033 [−0.039, −0.027] + | −0.033 [−0.038, −0.029] + | −0.030 [−0.034, −0.027] + |
| Kernel memory minus MLP | MSE | −0.084 [−0.224, 0.060] ∘ | 0.149 [0.059, 0.263] ▲ | −0.038 [−0.077, 0.010] ∘ |
|  | Direction error | −0.023 [−0.028, −0.017] + | −0.005 [−0.008, −0.002] + | −0.010 [−0.012, −0.009] + |
| BioMeme minus same-head MLP | MSE | −0.271 [−0.398, −0.165] + | 0.079 [0.019, 0.156] ∘<sup>†</sup> | 0.012 [−0.061, 0.134] ∘ |
|  | Direction error | −0.021 [−0.026, −0.016] + | −0.006 [−0.009, −0.003] + | −0.007 [−0.008, −0.005] + |
| BioMeme minus reference-informed MLP | MSE | 0.047 [−0.063, 0.152] ∘ | 0.109 [0.052, 0.181] ▲ | 0.882 [0.600, 1.332] ▲ |
|  | Direction error | −0.001 [−0.007, 0.005] ∘ | 0.001 [−0.003, 0.006] ∘ | 0.064 [0.059, 0.070] ▲ |
| Activity: BioMeme minus kernel memory | AUROC | 0.101 [0.075, 0.130] + | — | — |
| Cross-cell/same-cell correlation ratio | Ratio | 0.489 [0.336, 0.620] + | — | — |
| Compound transferability | *r<sub>s</sub>* | 0.362 [0.266, 0.450] + | — | — |
| RNA gain: high minus low TAS | Fraction | 0.028 [< 0.001, 0.058] ∘ | — | — |
| Neighbour enrichment: RNA minus RNA-key-block removal | Fraction | 0.015 [0.005, 0.025] + | — | — |
| Neighbour enrichment: RNA minus shuffled RNA | Fraction | 0.033 [0.020, 0.046] + | — | — |
| Transfer gain: strong minus weak | Fraction | 0.268 [0.216, 0.319] + | — | — |
| Kernel memory minus linear | MRR | 0.040 [0.027, 0.053] + | — | — |
| Kernel memory minus linear | Margin | 0.045 [0.039, 0.051] + | — | — |
| Kernel memory minus MLP | MRR | 0.034 [0.023, 0.046] + | — | — |
| Kernel memory minus MLP | Margin | 0.032 [0.027, 0.037] + | — | — |
| Kernel memory minus shuffled RNA | MSE | −0.280 [−0.368, −0.198] + | — | — |
| Kernel memory minus shuffled RNA | Direction error | −0.029 [−0.035, −0.023] + | — | — |
| Kernel memory minus RNA-key-block removal | MSE | −0.072 [−0.156, 0.012] ∘ | — | — |
| Kernel memory minus RNA-key-block removal | Direction error | −0.008 [−0.014, −0.002] + | — | — |
| BioMeme minus same-head MLP: low coverage | MSE | −0.500 [−0.837, −0.212] + | — | — |
| Low minus high coverage advantage | MSE | −0.400 [−0.746, −0.106] + | — | — |

Compounds tended to transfer to both target cells. After adjustment for A549 response covariates, transfer coefficients (target-cell changes projected onto A549 changes) agreed across MCF7 and U2OS (partial Spearman correlation 0.362 [0.266, 0.450]; 289 compounds; adjusted *P* = 0.0002). In an additional analysis using a reference cell line profiled in another batch, agreement was 0.503 (0.314–0.647; 80 compounds at 48 h). In this common population, the same-batch reference gave 0.252 across both times and 0.152 (*−*0.046 to 0.351) at 48 h. Further adjustment for both target-cell response strengths retained a positive correlation of 0.365 (0.105–0.571), consistent with compound-specific agreement beyond response strength (Supplementary Note 9.6). Transfer coefficients were higher for dopamine-receptor antagonists, selective serotonin-reuptake inhibitors, oestrogen-receptor antagonists and PARP inhibitors than for opioid antagonists and retinoid or vitamin D receptor agonists (Supplementary Table 69).

Shared responses supported prediction across further cell-line pairs. In LINCS morphology, reference transfer improved all 16 cell-pair–layout settings across six directed pairs (the two targeting MCF7 used development data), with pooled MSE reductions of 16.4–20.8% (Fig. 2d; Supplementary Data 4). In the 101-compound MCF7 replication cohort with complete data for both reference cell lines, A549 and U2OS reduced compound-averaged MSE by 24.0% and 23.8%, respectively. Transfer also reduced MSE in all six sci-Plex3 pairs, by 0.3–6.4% (Supplementary Fig. 3 and Supplementary Table 10).

Same-well Tahoe scoring covered 11 cell lines and 110 directed pairs (261 drugs; Fig. 2g; independent-well scoring of A549-referenced predictions in Fig. 3h). MSE decreased in every pair, with intervals excluding zero in 108. Direction error decreased with intervals excluding zero in all 110. Every reference line reduced same-well target error; A549 ranked ninth by median MSE reduction across its 10 targets.

**Fig. 3.**
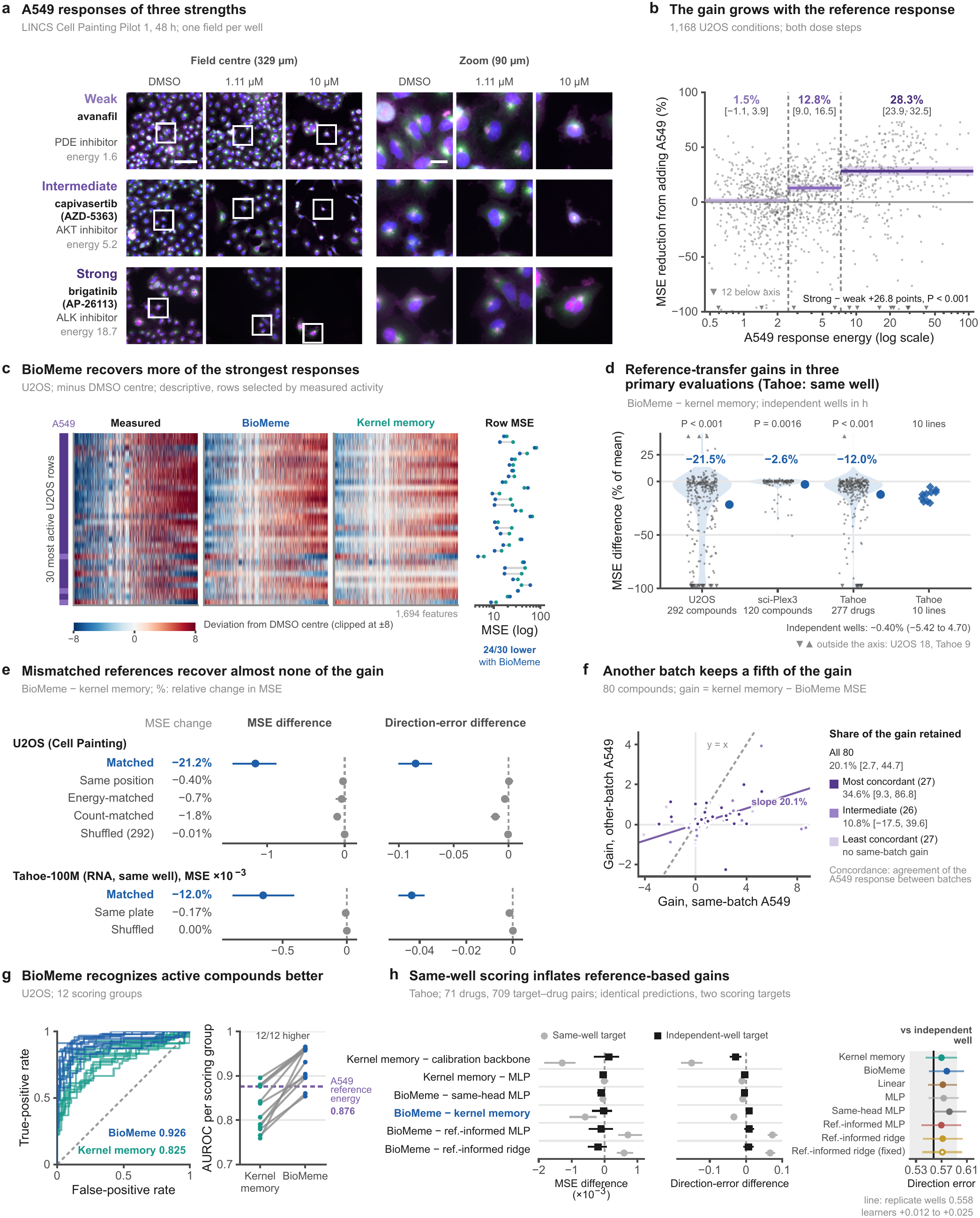
| Matched reference measurements reveal compound-specific responses. **a,** Preregistered Batch 2 A549 energy strata illustrated by Pilot 1 images (48 h); analysed profiles are Batch 2. DNA, blue; ER, green; AGP, magenta. Display ranges: Supplementary Table 28; field/zoom scale bars, 100/20 µm. **b,** Reference energy versus MSE reduction, 1,168 U2OS conditions; pooled gains and intervals within registered strata. Triangles indicate 12 conditions below *−*100%. The strong-minus-weak contrast uses a one-sided whole-compound bootstrap test (biological family H5). **c,** Thirty most active U2OS profiles (18 compounds; descriptive selection); features follow the first principal-component loading. The A549 bar marks energy strata. Colours saturate at *±*8; BioMeme lowers MSE in 24 profiles. **d,** Transfer gains in three screens and ten Tahoe targets, normalized to mean kernel-memory MSE; Tahoe uses same-well scoring. Triangles indicate 18 U2OS and nine Tahoe off-axis values. Displayed transfer *P* values use two-sided centred whole-compound bootstrap tests. **e,** Matched and mismatched references. U2OS comparisons use 241 compounds, except the shuffled control (292); same-position controls preserve well position. Wrong Tahoe compounds preserve plate. Energy- and count-matched wrong compounds were selected using A549 measurements only, and the transfer head was refitted. **f,** Cross-batch A549 transfer, 80 compounds. **g,** Activity area under the receiver operating characteristic curve (AUROC) in 12 groups; dashed line, reference-energy baseline. **h,** Identical Tahoe predictions scored against same- and independent-well targets: 71 drugs, 709 target–drug pairs. Negative contrasts favour the first model. Inset: direction errors versus the empirical replicate-well reference (post hoc; Supplementary Tables 29 and 49). Intervals are 95% whole-compound bootstrap intervals (10,000 draws); displayed *P* values are Holm-adjusted within their registered families. Source data are provided with this paper.

### Same-batch reference measurements carry compound-specific information

Adding A549’s response reduced U2OS MSE by 21.5%, from 5.776 for kernel memory to 4.534 for BioMeme (Fig. 3d; Table 1). The paired difference was *−*1.242 (*−*1.512 to *−*0.991), and direction error fell by 0.091 (0.077–0.106). In the 258 compounds and 827 conditions without low cell counts, transfer lowered MSE by 17.0% (difference 0.496, 0.335–0.691) and direction error by 0.060 (0.045–0.076); BioMeme still outperformed the same-head MLP on both endpoints (Supplementary Note 10.8). Reference response strength was measured as energy, the mean squared feature change between doses (Methods). Gains rose with reference-cell response strength, from 1.5% to 12.8% and 28.3% lower MSE across weak, intermediate and strong strata (Fig. 3a–c). The full 102-compound MCF7 replication showed a 25.2% MSE reduction and a 0.117 direction-error reduction (Supplementary Fig. 14b; Supplementary Table 26).

Low-cell-count conditions contributed 77.2% (69.7–83.5%) of the signed MSE gain, yet the full gain required the matched compound. Across 241 compounds, a wrong reference compound with the closest A549 cell-count trajectory recovered 8.3% (4.3– 12.7%) of the matched-compound MSE gain. Cell-count change and the first response principal component, each fitted as the sole reference input, recovered 10.3% and 25.7% of the full transfer gain (Supplementary Table 68). Removing the training-derived target principal component left BioMeme more accurate than kernel memory and the same-head MLP on both endpoints. Transfer increased MSE for 36.4% of conditions and 20.9% of compounds after averaging their four conditions (Supplementary Table 69).

Reference responses recovered higher-dose morphology when lower-dose transcription was weak. Among 84 compounds with low transcriptional activity scores (TAS), 43 were morphologically active at 24 h and 47 at 48 h, using a control-based threshold (Supplementary Fig. 24; Supplementary Table 31). At 24 h, activity occurred in 46 of 95 intermediate-TAS and 94 of 113 high-TAS compounds. Seven of the 43 low-TAS compounds were already active at the source dose. At the higher dose, 38 left the low-TAS stratum in the L1000 gene-expression assay, as did all 10 MCF7 replication compounds (Supplementary Note 9.3). For active low-TAS compounds at 24 h, between-compound *R*^2^ was 0.10 with the backbone, 0.09 with morphology-keyed memory and 0.08 with RNA added. Reference transfer raised it to 0.42 and, at 48 h, from 0.11 to 0.35. The reference-informed MLP reached 0.41 and 0.40; MCF7 replication showed the same benefit (Supplementary Table 21).

Reference transfer improved both transcriptomic evaluations. In sci-Plex3, using LINCS-derived kernel parameters, it reduced MSE by 2.6% and direction error by 0.010; both passed Holm correction. Replicate noise accounted for about 87% of kernel-memory MSE; transfer removed 19.8% of estimated reducible error (Supplementary Note 8; Supplementary Table 8). In Tahoe, MSE fell by 12.0% and direction error by 0.043 across 10 lines sharing wells with A549 (Fig. 3d). All 10 MSE estimates favoured BioMeme, eight intervals excluded zero, and all 10 direction intervals excluded zero (Supplementary Fig. 2; Supplementary Table 9). Tahoe replicate differences correlated more for the same drug than for other drugs on those plates (*r* = 0.355 versus 0.113). With A549 and U2OS on separate LINCS plates, the corresponding excess correlation was 0.008 (*−*0.001 to 0.017; Supplementary Note 9.9). Against independent wells, unchanged predictions for 71 Tahoe drugs retained memory’s direction advantage over the backbone and MLP, and BioMeme’s over the same-head MLP. Memory’s direction advantage over linear regression and MSE advantage over the backbone disappeared. Transfer no longer significantly reduced MSE, and it increased direction error; reference-informed MLP and cross-validated ridge lost their significant advantages over BioMeme (Fig. 3h). The eight displayed models had direction errors 0.012–0.025 above the replicate value of 0.558; fixed-penalty ridge had the lowest MSE (Supplementary Tables 29 and 49).

References were most useful when compound and batch matched. In 241 U2OS compounds, the matched reference reduced MSE by 21.2%, compared with 0.40% for another compound at the same well position. A wrong compound matched on mean reference-response energy reduced MSE by 0.7% (*−*0.6 to 2.0%; Supplementary Note 9.11). Matching on the four-condition energy trajectory recovered 8.5% (0.7– 15.9%) of the matched-compound gain (Supplementary Table 68). Shuffling identities contributed essentially no benefit, and wrong-compound Tahoe references from the same plate groups recovered 1.4% of the matched-compound MSE gain (Fig. 3e). A reference cell line from another batch retained 20.1% (2.7–44.7%) of the same-batch gain. Compounds with the most concordant responses between batches accounted for 84% of this retained benefit (Fig. 3f; Supplementary Table 18). Against a fifth U2OS replicate plate from the same batch, whose index was absent from A549 inputs, unchanged predictions retained 94.1% (85.1–104.9%) of their original-target MSE gain over kernel memory among 194 eligible compounds (Supplementary Note 9.10).

BioMeme improved active-treatment identification. U2OS activity discrimination (area under the receiver operating characteristic curve, AUROC) rose from 0.825 for kernel memory to 0.926 for BioMeme, improving in all 12 scoring groups (Fig. 3g; Supplementary Fig. 6). Reference-cell response energy alone gave 0.876, below BioMeme by 0.050 (0.020–0.081; additional analysis; Supplementary Note 9.7). In sci-Plex3, BioMeme reached 0.759, exceeding kernel memory by 0.065 and the same-head MLP by 0.036 (Supplementary Fig. 14c; Supplementary Table 13).

### Memory predicts response direction better than the learners it replaces

Memory predicted response direction more accurately in all nine preregistered comparisons (Fig. 4b; Table 1): BioMeme versus the same-head MLP, and kernel memory versus linear and neural learners, in each of three screens. All passed within-screen Holm correction, with direction advantages of 0.6–7.6% of comparator error. Tahoe used the registered tuned kernel. With the registered fixed kernel, Tahoe kernel memory had higher error than the MLP on both endpoints, whereas BioMeme retained lower direction error than the same-head MLP (Supplementary Table 9). MSE differences favoured memory in three of the nine comparisons, were not significant in five and favoured the MLP in one (Table 1). Training-compound selection of the linear penalty preserved memory’s direction advantage in U2OS, reserved MCF7 and sci-Plex3; MSE favoured memory in the two morphology populations but the selected-penalty linear learner in sci-Plex3 (Supplementary Table 65).

**Fig. 4.**
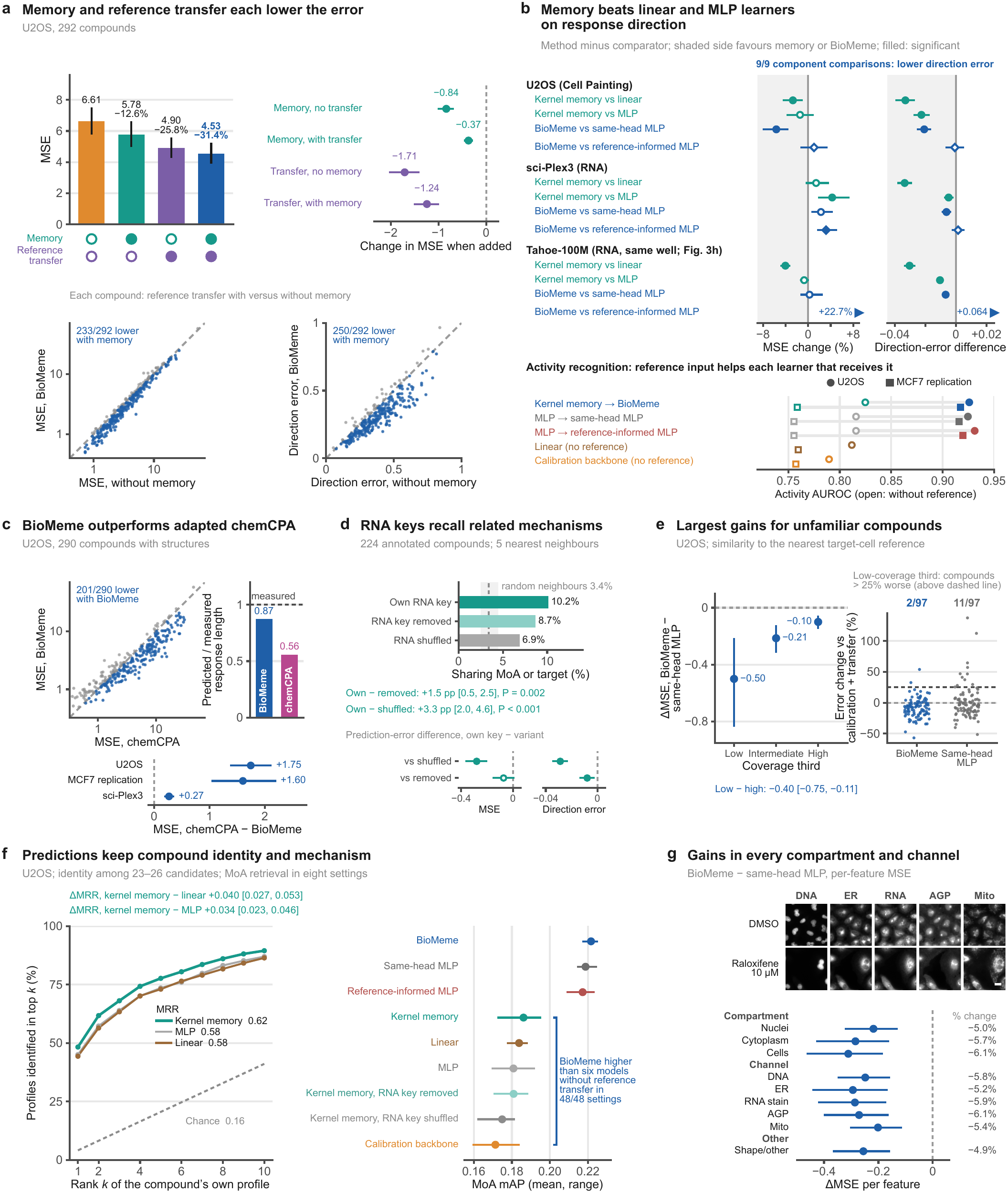
| Response memory improves learned-layer predictions and retains biological information. **a,** Memory–transfer ablation in 292 U2OS compounds: mean MSE, incremental component effects and BioMeme versus calibration-plus-transfer scatterplots. **b,** Learner replacements in U2OS (292 compounds), sci-Plex3 (120) and Tahoe (277). The nine component comparisons are memory versus linear or neural learners and BioMeme versus the same-head MLP in each screen. All nine favour memory for direction. Diamonds compare BioMeme with the reference-informed MLP, which reads reference responses directly; edge arrows mark off-axis Tahoe contrasts. Below, descriptive activity AUROC in U2OS and MCF7 replication; open/filled symbols denote absent/present reference information. **c,** Adapted chemCPA versus BioMeme in 290 U2OS compounds with structures. Response length is the predicted/observed profile Euclidean-norm ratio, without additional centring. Positive chemCPA-minus-BioMeme contrasts favour BioMeme. **d,** Mechanism or target sharing among five nearest neighbours for 224 annotated compounds. Random baseline: mean and central 95% range of 1,000 random-neighbour draws. Forests compare intact, removed and shuffled RNA key blocks. **e,** Coverage thirds (389, 390 and 389 conditions) defined by nearest-reference kernel affinity within prediction blocks. The strip shows the 97 compounds in the lowest third of mean condition-level coverage percentiles; the dashed line marks errors 25% above calibration plus transfer. **f,** Compound identification: mean reciprocal rank (MRR) among 23–26 candidates. Mechanism-of-action (MoA) retrieval: mean average precision (mAP), with means and ranges across eight settings; 48 comparisons concern six models without transfer. **g,** U2OS BioMeme-minus-same-head-MLP errors by compartment and channel. Example tiles show the five Cell Painting channels in A549 Pilot 1 fields at 48 h (DMSO, top; raloxifene 10 µM, bottom; RNA, nucleoli and cytoplasmic RNA; Mito, mitochondria; scale bar, 20 µm; display ranges, Supplementary Table 28). Intervals are 95% whole-compound bootstrap intervals (10,000 draws). Filled/open forest markers indicate significant/non-significant Holm decisions. Learner tests (**b**) and RNA-ablation error tests (**d**) use two-sided centred bootstraps; neighbour-sharing tests (**d**, H4a/H4b) use one-sided bootstraps. Displayed *P* values are Holm-adjusted. Source data are provided with this paper.

Memory and transfer each improved morphology beyond the other (Fig. 4a). Backbone MSE of 6.612 fell to 5.776 with memory, 4.905 with transfer and 4.534 with both. Memory added to transfer lowered MSE by 0.370 (0.278–0.463); transfer added to memory lowered it by 1.242 (0.991–1.512). With transfer present, memory lowered MSE for 233 and direction error for 250 of 292 compounds. A single coefficient on the reference change recovered 46% of transfer gain; ridge supplied most of the remainder, with a small significant contribution from gating (Supplementary Tables 14 and 23).

BioMeme outperformed simple baselines and the same-head MLP in U2OS. MSE was 43.0% below the mean-response baseline (Table 2) and 23.6% below a linear learner without reference transfer (Supplementary Table 7). Using the same-head MLP increased MSE by 0.271 (0.165–0.398) and direction error by 0.021 (0.016–0.026). Against reference-informed ridge using the same inputs and an inner-cross-validated penalty, BioMeme had 5.5% lower MSE, a difference of *−*0.266 (*−*0.364 to *−*0.175), and 0.024 (0.019–0.030) lower direction error (Table 2; Supplementary Note 9.8).

**Table 2.**
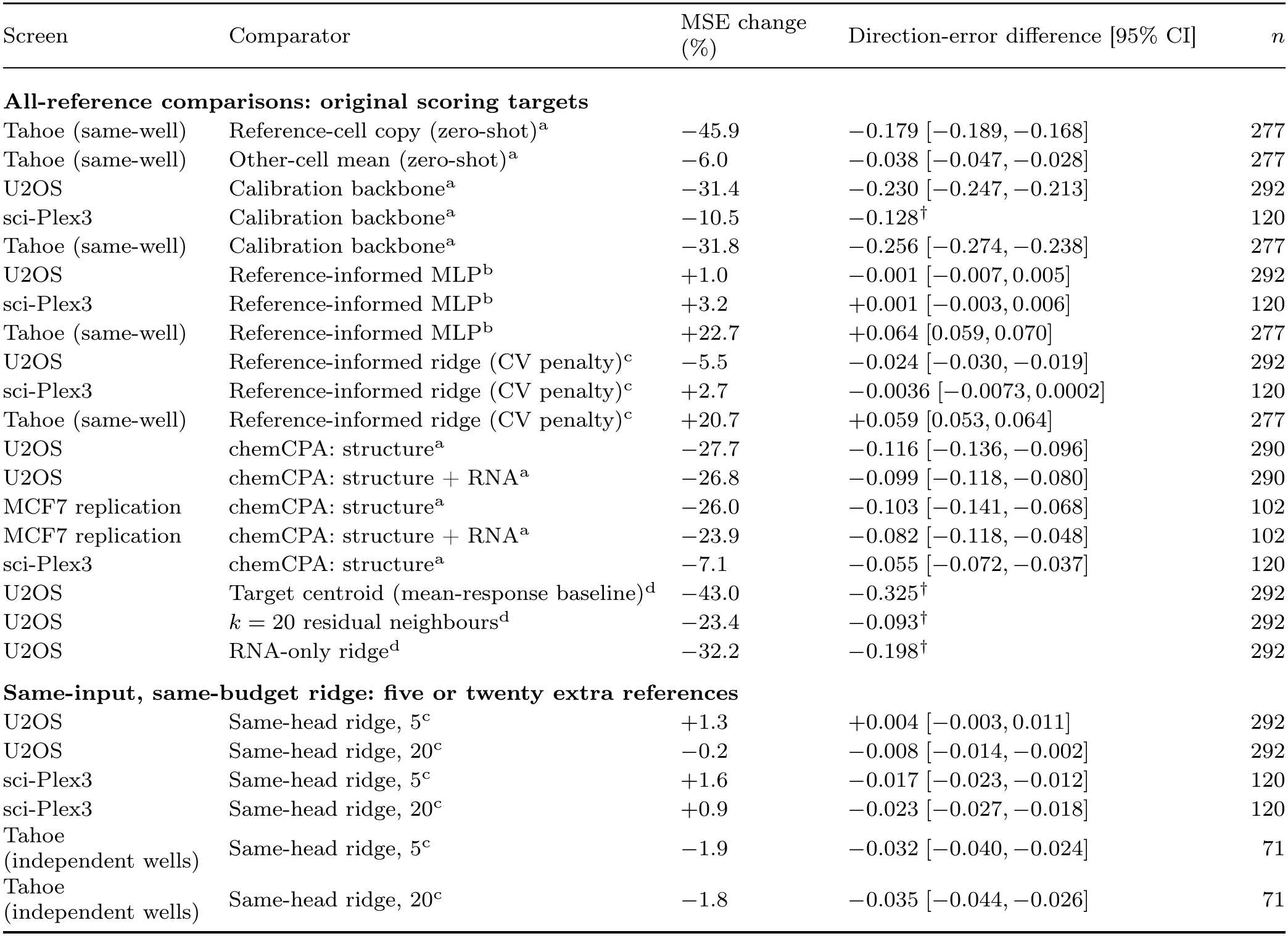
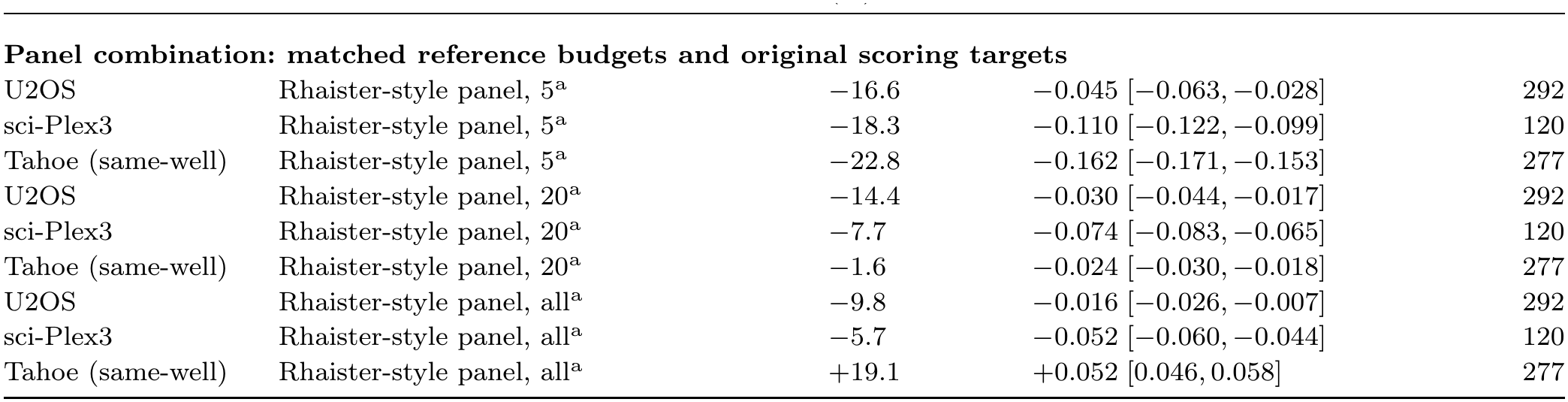
| Alternative prediction strategies and measurement budgets. **notes.** All differences are BioMeme minus comparator; negative values favour BioMeme. MSE change is 100[MSE_BioMeme_*/*MSE_comparator_ *−* 1], calculated from unrounded model means. Thus the U2OS ridge and target-centroid rows give the 5.5% and 43.0% reductions reported in the text. Direction intervals are paired whole-compound 95% intervals; four decimals are retained where rounding would obscure whether an interval excludes zero. CV denotes cross-validation. Superscripts identify the analysis: a, preregistered descriptive; b, confirmatory; c, additional; d, prespecified simple-baseline means whose paired comparisons against kernel memory were added after primary scoring. Additional analyses use nominal intervals. All-reference rows retain their original scoring targets and cohorts. Tahoe uses the registered tuned BioMeme and same-well scoring in this block. Fig. 3h and Supplementary Table 29 score these Tahoe predictions against independent replicate wells in 71 drugs. Small-budget rows compare the same five or twenty references beyond the fixed calibration panel. U2OS and sci-Plex3 retain their original targets; Tahoe uses fixed-kernel BioMeme and independent replicate targets for 71 drugs at the second dose step (709 target–drug pairs). Same-head ridge replaces memory with the linear residual learner and fits the transfer head to its own out-of-fold errors. With all references, reference-informed ridge receives the reference-informed MLP inputs and selects its penalty by inner cross-validation. Complete small-budget comparisons and curves are in Supplementary Tables 45 and 46. Panel rows compare the stated matched budgets; Tahoe uses the original 277-drug population, with fixed-kernel BioMeme at five and 20 references and the tuned model with all references. The Rhaister-style panel combination shares the calibration backbone, target-cell references and reference-cell measurements; BioMeme additionally uses query RNA in U2OS or the source-profile key in transcriptomic screens. Zero-shot comparators omit target-cell calibration/reference measurements. chemCPA receives the task’s reference-cell information and averages five predictions before scoring; U2OS uses the common 290-compound structure-available cohort and MCF7 the reserved 102-compound layout. *†*: only a mean difference is shown because the archived paired interval used another comparator. Supplementary Note 3 details inputs and adaptations.

BioMeme outperformed chemCPA^[21]^ adapted to receive the same reference measurements. Across 290 U2OS compounds, chemCPA had higher MSE by 1.75 (1.37–2.13); adding source-dose RNA left a difference of 1.67 (1.32–2.05), replicated in MCF7 (Fig. 4c; Table 2; Supplementary Fig. 10; Supplementary Table 22).

Reference input improved MLP activity discrimination in U2OS and MCF7 (Fig. 4b; Supplementary Table 13) and lowered U2OS MSE from 5.86 to 4.49. BioMeme’s full-panel U2OS accuracy matched the reference-informed MLP’s (MSE 4.53 versus 4.49; 95% interval for the difference, *−*1.4% to +3.4% of the MLP’s MSE; direction error not significantly different; Supplementary Table 7). Reserved MCF7 data replicated this result (MSE-difference 95% interval, *−*2.0% to +3.2% of MLP MSE; Holm *P* = 1.000). BioMeme’s MCF7 MSE advantage over the same-head MLP did not survive Holm correction (*P* = 0.064; Supplementary Table 26). In sci-Plex3, BioMeme’s MSE exceeded reference-informed MLP and cross-validated reference-informed ridge by 2.7–3.2%, with similar direction error; screen-specific kernel tuning removed the significant MSE differences (Supplementary Tables 8, 36 and 37). Both comparators were more accurate on both endpoints in same-well Tahoe, but their significant advantages disappeared against independent wells (Fig. 3h; Supplementary Table 29). An exploratory expanded transfer grid lowered Tahoe MSE from 4.76 to 4.03 (*×*10*^−^*^3^), a 15% reduction. Its excess over the reference-informed MLP narrowed from 22.7% to 3.9%, while the MLP retained lower direction error (Supplementary Note 8). With the registered tuned kernel in U2OS, the reference-informed MLP had lower error on both endpoints (Supplementary Table 7).

BioMeme fitted and predicted within seconds on one CPU thread. Across 12 U2OS folds with 20 references, its MSE was 2.2% higher than the reference-informed MLP’s (Supplementary Table 52). From prepared inputs, its workflow took 7.7–8.9 s per fold (median, 8.3 s), versus 57–229 min (median, 138 min) for neural tuning and fitting on the same CPU. Paired ratios were 420–546 in six folds needing no neural-search extension and 1,432–1,710 in six including prescribed extension and original-configuration sensitivity fits (Fig. 1f; Supplementary Table 58). Both used one thread per process; neural fits ran four concurrent workers. Supplementary Note 13 defines workflow boundaries. All 12 replays reproduced archived BioMeme predictions exactly.

### Memory weights favour related mechanisms and limit large errors on unfamiliar compounds

Measured transcription helped organize the memory (Fig. 4d; Supplementary Fig. 12). Among 224 annotated compounds, the five nearest neighbours shared a mechanism or drug target in 10.2% of cases, compared with 3.4% for random neighbours (additional analysis; Supplementary Note 10). Removing the RNA key reduced sharing to 8.7%, and shuffling RNA identities reduced it to 6.9%; both differences were significant (Table 1). Shuffling RNA also increased MSE by 0.280 and direction error by 0.029. Removing the RNA key, while retaining the calibration geometry fitted with RNA, increased direction error by 0.008 without significantly changing MSE.

All 1,168 U2OS predictions expanded through calibration, memory and both transfer terms into weighted sums of named measurements, reconstructing the archived predictions within 6*×*10*^−^*^14^ (Supplementary Fig. 17; Supplementary Table 67). Deleting the five or ten compounds with the largest absolute memory weights produced 4.5- or 3.7-fold larger root-mean-square prediction changes than random deletions, with greater MSE increases (Supplementary Table 67).

Memory’s measured-residual weights sum to one. Raltitrexed had the largest positive weight (0.10) for the prespecified pemetrexed example; both inhibit thymidylate synthase (Fig. 1c; Supplementary Fig. 23). Across 224 annotated compounds, the five largest positive weights shared mechanisms or targets in 11.2% (8.8–13.7%) of cases, a 3.3-fold enrichment over random selection. Sharing exceeded nearest-neighbour selection by 1.0 (0.2–2.0) percentage points (Supplementary Note 10.6). Kernel weights also exceeded a linear learner’s equivalent weights in annotation sharing (11.2% versus 9.8%, difference 1.4 [0.6, 2.2] percentage points) and concentrated on fewer compounds (median effective number, 68 versus 105; Supplementary Table 67).

Memory’s advantage over the same-head MLP increased where similar measured compounds were scarce: 0.50 MSE in the low-coverage third versus 0.10 in the high-coverage third; against the reference-informed MLP, the low-coverage difference was +0.024 (*−*0.260 to 0.290). Among the 97 low-coverage compounds, BioMeme made two errors exceeding the common calibration-plus-transfer baseline by more than 25%. The corresponding counts were 11 for the same-head MLP, three for the reference-informed MLP and five for reference-informed ridge (Fig. 4e; Supplementary Figs. 5 and 18; Supplementary Note 10). BioMeme’s advantage over reference-informed ridge was also greatest in the low-coverage third (0.52 versus 0.13 in the high-coverage third).

Memory predictions also preserved compound identity, ranking the correct compound higher among the candidates than a linear learner or an MLP (increases in mean reciprocal rank, MRR, of 0.040 [0.027, 0.053] and 0.034 [0.023, 0.046]; Fig. 4f; Supplementary Table 12). BioMeme retrieved mechanisms better than six models without reference transfer in all 48 comparisons (Fig. 4f; Supplementary Table 12). In centred within-prediction MoA retrieval, its mAP difference from the reference-informed MLP was 0.008 [*−*0.002, 0.018]. Against the same-head MLP, BioMeme improved all nine feature groups and had favourable estimates across all 29 prespecified strata (Fig. 4g; Supplementary Tables 14 and 15).

### Beyond a calibration panel, few target-cell references are needed

With 20 additional target-cell references, BioMeme retained at least 80% of the full-panel linear learner’s improvement in all three screens (Supplementary Fig. 25). These references supplemented the 60–64 calibration compounds. U2OS MSE was 4.844, compared with 4.534 for the full panel (Fig. 5b; Supplementary Table 16). Even without additional references, fixed-kernel BioMeme had lower error than full-panel fixed-kernel memory on both Tahoe endpoints, and in U2OS it had lower MSE but higher direction error. BioMeme had 45.9% lower Tahoe MSE than A549 copy and lower direction error than averaging other measured lines (Fig. 5f; Table 2). On the 71-drug independent-well subset, it still outperformed A549 copy on both endpoints; differences from the other-line mean were not significant (Supplementary Table 70).

**Fig. 5.**
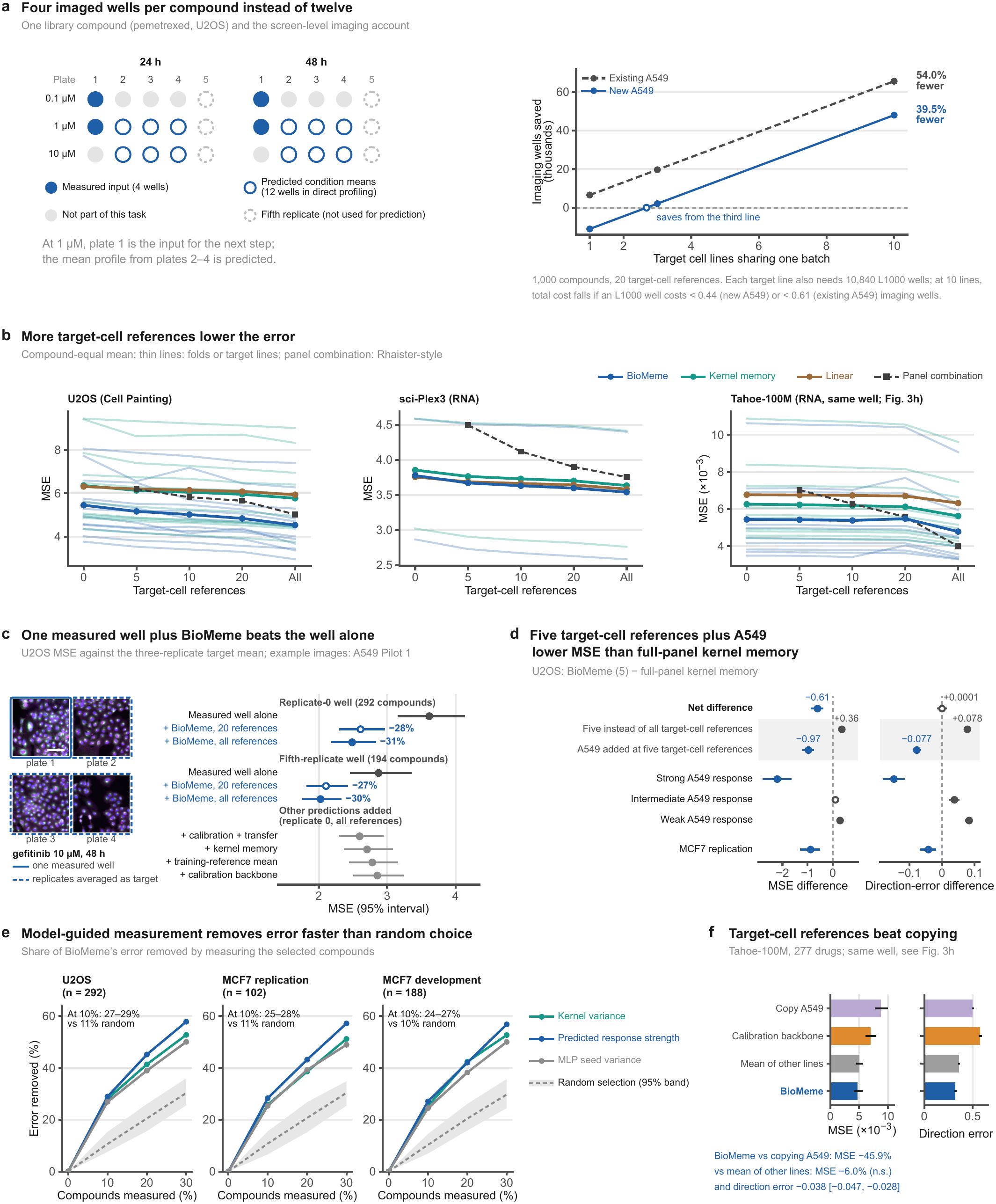
| Small reference panels support measurement prioritization, single-well estimation and reduced imaging. **a,** Four U2OS input wells per library compound predict four target-condition means, each otherwise measured in triplicate. Pemetrexed illustrates the design. The fifth replicate supplies the separate screening evaluation (Supplementary Note 12). The cost ledger covers 1,000 compounds, 20 references and 1–10 target lines sharing one batch (Supplementary Table 19). **b,** MSE versus reference budget. Tahoe uses fixed kernel parameters and linear penalty throughout. Thin curves: BioMeme and kernel-memory folds/target lines; dashed curves: Rhaister-style panel combination. The sequential 80%-retention test is in Supplementary Fig. 25. **c,** Single-well estimation in U2OS: one measured target-dose well alone or combined with BioMeme’s prediction (20 or all target-cell references), scored against the mean of replicates 1–3. Replicate-0 and fifth-replicate wells cover 292 and 194 compounds; percentages are MSE changes relative to the well alone. Other added predictions use the replicate-0 well and all references. Mixing coefficients were selected by nested cross-validation (Supplementary Notes 10.7 and 14.2; Supplementary Table 66A and B). Tiles show example fields from four replicate A549 Pilot 1 wells of gefitinib (10 µM, 48 h; solid frame, the measured well; dashed frames, replicates averaged as the target); they are not part of the U2OS scoring data. DNA, blue; ER, green; AGP, magenta; scale bar, 100 µm; display ranges, Supplementary Table 28. **d,** Five-reference BioMeme versus full-panel memory without transfer: MSE is 10.5% lower (5.17 versus 5.78); direction error differs by +0.0001, with an interval spanning zero. The joint one-sided bootstrap criterion (*α* = 0.025, both endpoints) was not met. Shaded rows decompose effects descriptively; filled markers denote intervals excluding zero. **e,** Error removed by measuring compounds ranked by kernel variance, MLP seed variance or predicted strength; random band, central 95% range of 100 selection orders. **f,** Tahoe predictions versus A549 copy or other-line means. Panels **b,f** use same-well Tahoe targets; independent-well comparisons are in Supplementary Tables 45–47 and 70. Unless specified, intervals are 95% whole-compound bootstraps (10,000 draws). Source data are provided with this paper.

With five or 20 target-cell references, BioMeme predicted direction more accurately than the Rhaister-style panel combination^[41]^ in all six screen–budget settings and had lower MSE in all six; the MSE interval included zero only at 20 Tahoe target-cell references (Tahoe scored in the same well; independent-well scoring in Supplementary Table 47). Across all nine settings, eight direction comparisons favoured BioMeme, which also used query RNA (U2OS) or the source-profile key (transcriptomic screens), absent from the panel combination (Table 2). With the full Tahoe panel, the panel combination was more accurate on both endpoints.

The memory learner itself contributed at small budgets. With the same key and budget, kernel memory predicted direction more accurately than a linear learner at five and 20 target-cell references in all three screens (Supplementary Table 45). Against reference-informed and same-head ridge, BioMeme improved direction without a significant MSE increase in two of four U2OS, two of four independent-well Tahoe and one of four sci-Plex3 comparisons; ridge had lower MSE in the other three sci-Plex3 comparisons (Supplementary Table 45).

A549 partly compensated for reducing the target-cell panel in morphology (Fig. 5d). With five target-cell references, U2OS BioMeme had 10.5% lower MSE than full-panel kernel memory, and MCF7 replication favoured BioMeme on both endpoints. Strong reference responses supported improvement on both endpoints, whereas weak responses did not offset the smaller panel (Supplementary Fig. 9; Supplementary Tables 17 and 26). The 20-reference MSE difference from the reference-informed MLP had a 95% interval of 0.02–4.68%; the direction-error difference was 0.002 (*−*0.005 to 0.009; Supplementary Table 52). Two stricter registered budget criteria were not met: retention against the reference-informed MLP, and joint improvement on both endpoints with five references (Supplementary Table 7).

Sharing a reference cell line reduced imaging wells relative to triplicate profiling (Fig. 5a). Each target line used four query wells in a 16-well design to predict four target means otherwise measured in 12 wells. Calibration and references added shared costs (Supplementary Table 19). For 1,000 compounds, 20 references and 10 target lines sharing one batch, a new A549 reference reduced imaging wells by 39.5%, recovering its cost by the third target line; an existing A549 reference reduced wells by 54.0%. Including 108,400 L1000 wells, total cost decreased if each cost less than 0.44 (new A549) or 0.61 (existing A549) imaging wells. Profiling each predicted condition once would use 40,480 imaging wells and no L1000 wells (Supplementary Table 19). U2OS between-compound *R*^2^ at this budget was 0.39 (full panel, 0.42; replicate reference, 0.89), and activity AUROC was 0.917 (full panel, 0.926; Supplementary Tables 39 and 41).

A morphology-keyed variant retained most of BioMeme’s full-panel accuracy (between-compound *R*^2^, 0.41 versus 0.42) while requiring RNA only for calibration compounds. Its separate full-panel cost scenario used 6,400 L1000 wells and reduced imaging wells by 32–47%; total cost fell below direct profiling when an L1000 well cost less than 6.1 or 9.0 imaging wells for a new or existing A549 reference cell line (Supplementary Table 19). Both ledgers charge source wells only to BioMeme.

Combining prediction with direct measurement improved response estimation. With weights fitted without queried compounds, adding BioMeme’s prediction to one target-dose well lowered MSE from 3.62 to 2.61 with 20 references and 2.49 with the full panel (Fig. 5c). On 194 compounds with fifth replicates, MSE fell from 2.87 to 2.11 and 2.03, reductions of 27–31% overall. At the intermediate dose, whose well is already imaged, MSE fell from 2.92 to 2.09 and 1.99 without an additional well. Across budgets and replicate evaluations, BioMeme combinations lowered MSE relative to calibration, memory and training-reference-mean combinations, and direction error relative to calibration plus transfer in all four settings, with lower MSE in three (Supplementary Table 66). The direct well alone was more accurate than prediction alone (Supplementary Note 10.7).

Model-based rankings identified measurements that removed prediction error ^[47]^. Kernel variance correlated with BioMeme’s error in U2OS (Spearman correlation 0.575) and MCF7 replication (0.423). After adjustment for predicted and measured response strengths, the correlations were *−*0.042 (*−*0.107 to 0.026) and *−*0.112 (*−*0.203 to *−*0.015; Supplementary Table 70). Retrospectively measuring the 10% of U2OS compounds with the largest kernel variance removed 28.0% of total error, versus 10.6% for random selection. Ranking by predicted response strength removed 28.9% and by MLP seed variance 26.9%, against 7.6–15.3% for random selection (central 95% range across 100 orders; Fig. 5e). These rankings reduced error but had lower active-compound recall than random selection (Supplementary Fig. 8 and Supplementary Table 20).

Each method chose 146 U2OS conditions for follow-up among those inactive at the lower dose, using 20 target-cell references. Choices were fixed before scoring on fifth replicate plates from the same screen. BioMeme selected 97 morphologically active conditions, compared with 97.8 for the reference-informed MLP (five-seed mean) and 99 for reference-energy ranking. Neither difference was significant (both Holm-adjusted *P* = 0.5448). Random selection expected 45.2 hits. Among secondary rankings, unfitted A549 activity at the target dose found 104 and kernel memory alone 74 (Supplementary Note 12 and Supplementary Fig. 21).

## Discussion

Biological response memory replaces a learned response layer with experiments that a virtual cell can recall and inspect. Memory predicted direction better in all nine preregistered learner comparisons, and full-panel U2OS accuracy matched a network reading reference responses directly. Predictions remained traceable to measurements, with refitting in seconds on one CPU thread. The query’s lower-dose state links target-cell memory to same-compound transfer, separating each route’s evidence.

Measured-state keys grouped biologically related experiments. Shuffling RNA increased error and reduced mechanism or target sharing (Fig. 4d), and the largest positive weights favoured shared mechanisms. Structure-based and raw-RNA keys gave accuracy similar to the RNA-response key, and the structure key had higher mechanism sharing (11.9% versus 10.2%). Laboratories can use available descriptions (Supplementary Tables 23 and 69). Solving for memory weights improved on averaging 20 neighbours under the same key (MSE difference, *−*0.141 [*−*0.226, *−*0.059]). The Gaussian-process correction returns to the mean residual far from measured examples^[48–51]^, while reference transfer remains available. This behaviour is consistent with the larger advantage over the same-head MLP for unfamiliar compounds (Fig. 4e).

Transfer identifies useful measurement designs. References should profile the same compound, batch^[52,53]^ and doses. Reference-cell responses recovered higher-dose morphology that weak lower-dose transcription forecast poorly ^[54]^. In sci-Plex3, transfer between adherent A549 and MCF7 lines (3.0–6.4%) exceeded transfer involving suspension K562 cells (0.3–1.5%), suggesting a role for cell-type similarity (Supplementary Fig. 3). Most LINCS transfer gain arose in low-cell-count conditions. Cell-count change and the leading response component recovered 10.3% and 25.7% of the gain. Wrong compounds with the closest cell-count trajectories recovered 8.3%, supporting compound-specific information beyond measured cell abundance. Shared within-well variation inflated Tahoe gains, requiring independent-well evaluation ^[55,56]^. These findings complement evidence that spreading cells across more conditions improves generative prediction^[18]^. Next-dose activity prediction could inform Cell Painting toxicology ^[45]^.

State-dependent recall connects BioMeme to neural memory and retrieval, and reference transfer to panel regression and in-context reuse^[36,39–41,57–62]^. At 20 references, its fitting workflow was at least 420 times faster than the input-matched MLP’s on the same CPU. Local refitting makes response memory accessible without GPU infrastructure. Exact kernel solves scale cubically, motivating Nyström features, inducing points or rank-one updates for larger memories^[48]^. GENESIS^[18]^ or pretrained representations^[63–66]^ could supply backbones or keys.

Predictions extended one adjacent-dose step, and cross-batch references retained about a fifth of transfer gain. The reference-informed MLP had 2–4% lower U2OS MSE with five or 20 references (Supplementary Table 52). Three small-budget sci-Plex3 comparisons favoured ridge on MSE despite lower or similar BioMeme direction error. In sci-Plex3, kernel tuning removed the significant full-panel MSE differences. Unfitted A549 activity found at least as many lower-dose-inactive conditions becoming active as BioMeme, making it a strong first filter. In BBBC021 ^[67]^, kernel memory lowered the backbone’s MSE in the one-step task but not in the two-step task, and did not differ significantly from linear or neural learners (Supplementary Note 6).

The supplied interface supports fitting, tracing and updating predictions, with prepared tasks for benchmarking (Supplementary Data 42). Prospective screens, distant cell types, genetic perturbations and time courses offer extensions. Robot scientists and AI agents already plan and run experiments^[32,33]^. An automated laboratory could test memory-guided screening in a closed loop, using predicted response strength to select measurements and linear solves to incorporate new responses. Treating experiments as memory makes every measurement, whether made by hand or by robot, evidence for a virtual cell’s next prediction.

## Methods

### Study design and data

Primary comparisons used LINCS Cell Painting, sci-Plex3 and Tahoe-100M, with A549 as the reference cell line. Permitted reference-cell inputs spanned its complete dose–time screen (Supplementary Table 44).

Each task assigned whole compounds to calibration, target-cell reference or query roles. Calibration compounds fitted preprocessing and the source-to-target backbone. Additional target-cell references numbered 0, 5, 10, 20 or all available compounds; calibration compounds remained at zero references. All observations of each compound shared fold assignments (Supplementary Fig. 1). Query targets were reserved for scoring. Supplementary Tables 1–3 list populations, permitted inputs, hyperparameters and registrations. Supplementary Note 6 describes the separate BBBC021 evaluation.

#### LINCS Cell Painting and L1000

The LINCS Cell Painting Batch 2 resource, cpg0004-lincs^[46]^, profiles A549, MCF7 and U2OS in layouts ASG003, LKCP001 and LKCP002. Cell Painting measures morphology across fluorescent channels^[7]^. Profiles were available at 24 and 48 h. Among the first four plates, replicate 0 supplied inputs and the mean of replicates 1–3 defined targets. Two adjacent transitions, lower to intermediate and intermediate to higher dose, gave four rows per compound across both times. Recorded nominal concentrations determined dose covariates; Pilot 1 and Batch 2 dose labels differed slightly, and final Batch 2 assay concentrations were not independently verified (Supplementary Note 1). Earlier-time and lower-dose inputs followed Supplementary Table 44.

We retained 1,694 morphological features shared with independent A549 Pilot 1. Official profiles were normalized to medians and median absolute deviations (MADs) of same-plate wells labelled dimethyl sulfoxide (DMSO). On the 26 ASG003 plates (24 task plates and two fifth-replicate plates), this label comprised six solvent and 12 zero-volume blank wells (Supplementary Note 10.9; Supplementary Table 63). Without further normalization, non-finite entries became zero and values were clipped to [*−*10, 10]. Matched 24-h L1000 signatures supplied 978 landmark genes^[4,54]^. Source-dose RNA informed morphology at 24 and 48 h; the second transition used intermediate-dose RNA.

U2OS confirmation comprised 292 query compounds and 1,168 rows across three layout directions. ASG003 to LKCP002, LKCP002 to ASG003 and ASG003 to LKCP001 contained 93/64, 98/60 and 101/64 query/calibration compounds, respectively. Four whole-compound query folds defined each direction’s reference pools. Source calibration and target-cell reference layouts were distinct. Supplementary Note 1 specifies the separate MCF7 development and reserved-layout replication populations.

#### sci-Plex3

sci-Plex3 profiles chemical perturbations in A549, MCF7 and K562 by multiplexed single-cell RNA sequencing ^[5]^. We used 24-h measurements at four doses, giving three adjacent transitions. Replicate 1 supplied inputs and replicate 2 supplied targets, from distinct treated wells on different hash plates. A549 reference changes used next-dose replicate 2 minus lower-dose replicate 1; sequencing-library pool identities were unavailable. Conditions required at least 20 cells per replicate; compounds required all four doses in reference and target lines. A fixed 60-compound calibration panel came from 175 compounds eligible in all three lines. Queries comprised 117 compounds in MCF7 and 118 in K562, with a union of 120.

Condition–replicate counts were summed, normalized to counts per million human-gene transcripts and transformed by log(1 + *x*). The landmark filter required mean solvent-cell expression of at least one count per million in every line, retaining 913 of the 978 L1000 landmark genes. Disjoint, metadata-defined solvent-cell subsets supplied reference pseudobulks. Each gene was centred on its median across those pseudobulks and divided by the larger of 1.4826 *×* MAD and 0.05. Values were clipped to [*−*10, 10].

#### Łahoe-100M

Tahoe-100M profiles mixed-cell-line perturbations at 24 h^[6]^. We used pseudobulk counts from chem-perturbridge, the fallback data product specified in the registered plan. A seeded, metadata-only round-robin selected 10 target lines from 10 tissues, with A549 references. Transitions were 0.05 *→* 0.5 and 0.5 *→* 5 µM. Conditions required at least 50 treated and 50 same-plate DMSO cells; compounds required all three doses. A 60-compound calibration panel and four global compound folds yielded 277 distinct queries across targets.

Treatment and same-plate DMSO counts were separately summed, normalized to 10^4^ counts across all input genes and log(1 + *x*)-transformed; their differences defined responses. A DMSO-only filter retained 928 landmark genes. Designed pools contained 50 cell lines; each plate had 94 treatment wells and two DMSO wells, pooled within cell line for controls. Different doses occupied different plates and used their respective controls. These log-fold changes differ from other resources’ robust z-scores, so absolute MSEs were compared within resource. Supplementary Note 1 gives preprocessing details.

### BioMeme

#### Calibration backbone

The backbone estimated the target profile from permitted source-profile streams *x_ℓ_*, *ℓ* = 1*, . . . , L*:

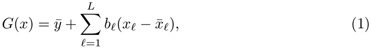

where *ȳ* and *x̄_ℓ_* are the target and source-stream means and *b_ℓ_* is one scalar slope per stream, shared across output features. Compound-weighted least squares fitted means and slopes on calibration compounds, using a minimum-norm solution for collinear streams. Eight whole-compound folds generated calibration residuals. Target-cell reference residuals used the backbone fitted on all calibration compounds. The memory stored the resulting residual responses.

#### Kernel memory

Memory keys described measured source states and dose transitions. Morphology keys contained 16 source-profile principal components, 16 RNA-derived response components and 7 log-dose/time covariates. The RNA response map used 887 independent A549 Pilot 1 compounds (1,772 dose tasks), excluding Batch 2 identities by identifier or first InChIKey block. Eight ridge providers predicted cross-fitted morphology residuals, each excluding one of eight Pilot 1 compound groups. This external map was fixed before confirmation. Within each direction, calibration compounds fitted feature standardization, weighted principal components and block root-mean-square scaling. Transcriptomic keys contained 16 source-profile components and standardized log source and target doses, giving 18 coordinates.

For key coordinates *z_i_*and *z_j_*, the kernel was

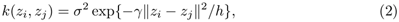

where *γ* sets the kernel width, *h* is the median non-diagonal squared distance between calibration coordinates and *σ*^2^ is their mean squared norm. Let *K* = *k*(*Z, Z*) for training coordinates *Z*, with *n* rows and residual matrix *R*. Row weights *w_i_* sum to one and give each compound equal total weight; *W* = *n* diag(*w*). The residual intercept *r̄* = *w^⊤^R* was fixed before fitting. With ridge penalty *λ*, the coefficient matrix *A* minimized

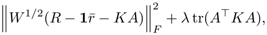

giving

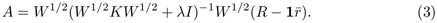

For a query key *z*, the memory prediction was *ŷ*_memory_ = *G*(*x*) + *M*, with memory term *M* = *r̄* + *k*(*z, Z*)*A*. This kernel-ridge fit ^[48,49]^ was repeated for every fold, provider and reference budget. Equation 3 gives the Gaussian-process posterior-mean coefficients for constant mean *r̄* and noise variance *λ/W_ii_*; Eq. 4 is the equivalent-kernel form^[68,69]^. Linear smoothers, including ridge, admit this decomposition; the kernel makes weights depend on measured-state similarity.

For one provider, let *s_z_* = *k*(*z, Z*)*W* ^1*/*2^(*W* ^1*/*2^*KW* ^1*/*2^ + *λI*)*^−^*^1^*W* ^1*/*2^. The memory contribution uses stored experiments:

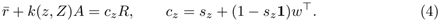

Entries of *c_z_* sum to one but can be negative, unlike kernel similarities. Averaging provider-specific contributions gives the final memory term. Coefficients retain experiment identifiers and exactly reconstruct the memory term (Supplementary Note 10).

#### Gated transfer head

The transfer head added the same compound’s reference-cell response. Let *D* be its measured change over the query dose transition and *e* = ‖*D‖*^2^*/p*, with output dimension *p*. BioMeme predicted

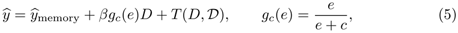

with *g_c_*(*e*) = 1 for all *e* when *c* = 0. The trajectory *D* concatenates the compound’s reference-cell dose changes. We fitted *β* without an intercept; weighted principal-component ridge regression fitted the remaining out-of-fold error through *T* , whose inputs were *D* and a centred, normalized *D*. Selected ranks and gates are listed in Supplementary Table 43. Response strength gated only the direct-transfer term, *B* = *βg_c_*(*e*)*D*; the ridge correction *T* was ungated.

#### Parameter selection

The fixed *γ* = 1, *λ* = 100, 39-coordinate morphology key and eight-provider construction came from A549 development on ASG003-to-LKCP002 and reverse directions, with 64 and 60 fitting compounds. These choices were informed by previously explored A549 outcomes and locked before U2OS confirmation opened. Held-out U2OS next-dose responses did not select them. Subsequent MCF7 development informed the transfer design and coverage hypotheses (Supplementary Note 1). LINCS and sci-Plex3 retained the fixed kernel parameters. Tahoe selected them by eight-fold training-compound cross-validation at full budget and used fixed values below it. Fixed-kernel Tahoe and tuned-kernel LINCS and sci-Plex3 variants were reported separately (Supplementary Tables 7–9 and 26; Supplementary Fig. 11b,c).

The transfer head fitted the base learner’s own out-of-fold errors. Eight-fold training-compound cross-validation selected *c ∈ {*0, 1, 2, 4, 8, 16, 32*}*, rank *∈ {*16, 32, 64, 128, 256, full*}* and ridge penalty *λ_T_ ∈ {*10^3^*, . . . ,* 10^8^*}*. Recorded grid order broke ties. Each head-selection fold refitted *β*. The selected head fitted all permitted training compounds. Formal analyses retained the fixed head grid. Kernel tuning used *γ ∈ {*0.25, 1, 4*}* and 11 *λ* values from 10 to 10^6^. One predefined boundary extension was allowed if it strictly reduced training-fold error. Supplementary Note 2 specifies centring, scaling and selection.

With fitted calibration coefficients, keys, gates and projections fixed, expanding the calibration means, memory residuals and transfer smoother gives a weighted sum of named measured profiles for the complete prediction. All 1,168 U2OS reconstructions agreed within 6 *×* 10*^−^*^14^ (Supplementary Note 14.3). Memory updates require a linear solve; complete-predictor updates repeat transfer-head selection.

### Reference selection and comparators

Greedy facility location selected target-cell references. Each added compound maximized the compound-weighted sum of query inputs’ largest kernel similarities to selected references. Lexicographic tie-breaking gave deterministic, nested reference sets shared by all models. Selection used permitted source inputs and excluded query targets. Tahoe used only the first transition’s source state. Each budget restricted residual fitting, training folds and transfer-head fitting separately.

The linear learner used compound-weighted ridge regression on the same coordinates, with an unpenalized intercept and a fixed penalty of 1,000 in LINCS and sci-Plex3; Tahoe selected its all-reference penalty on the kernel penalty grid. The fixed penalty of 1,000 had been selected by eight-fold training-compound cross-validation in both A549 development directions on 8 October 2026. The MLP had one hidden layer of 256 Gaussian error linear units (GELUs) and a zero-initialized output layer. Full-batch AdamW used learning rate 0.001. Eight-fold compound cross-validation selected weight decay and epoch count. When a boundary was selected, predefined rules extended the epoch limit from 400 to 800 or added weight decay 1. Final fits used five fixed seeds. The same-head MLP replaced memory and fitted the same transfer-head architecture to its own out-of-fold errors. Same-head ridge applied this construction to the linear learner. The reference-informed MLP added reference-response and trajectory principal components as inputs, as in BioPert ^[42]^, and fitted its own direct-transfer coefficient. Reference-informed ridge used the same inputs and selected its penalty by inner cross-validation.

Other comparators were the backbone, target and residual centroids, 20-neighbour residual retrieval and RNA-only ridge. The Rhaister-style panel combination^[41]^ learned ridge weights from A549 trajectories to combine the same target-cell reference residuals. Penalty selection left out one reference compound at a time. The panel combination shared BioMeme’s backbone and reference measurements. BioMeme additionally used query RNA in U2OS, or query source-profile coordinates in the transcriptomic resources. Adapted chemCPA^[21]^ received the task’s source and reference measurements. U2OS used structure-only and structure-plus-RNA versions; sci-Plex3 used structure. Tahoe also included reference-cell copying and averages across eligible non-target lines. Supplementary Note 3 specifies inputs, adaptations and aggregation. Pretrained State predicts perturbed populations from control cells and perturbation embeddings^[63]^, a different input setting from the treated, lower-dose query here. Tensor completion^[43]^ imputes whole drug–cell profiles under different input permissions and was not tested; Supplementary Table 38 compares related approaches’ inputs and evaluations.

### Evaluation and statistics

Simple and shuffled baselines and replicate references calibrated the metrics^[56]^; identification, mechanism retrieval and activity assessed compound-specific information (Supplementary Tables 12 and 13). Between-compound *R*^2^ (Supplementary Table 21) measures explained variation after removing each scoring group’s mean prediction error. Scoring groups contain compounds sharing context: layout, cell line, time and dose transition. Settings share compounds, so counts across settings are descriptive.

#### Endpoints

Primary row endpoints were feature-averaged MSE and direction error, 1 *−* cos(*y − o, y − o*), where *y* is the measured target profile. The fixed centre *o* was the condition-specific mean target profile of calibration compounds, shared across budgets. Rows were averaged within compound. Multi-target transcriptomic analyses then averaged available target lines within compound and weighted compounds equally. Neural losses were scored per seed before averaging. The registered main chemCPA readout scored the mean of five predictions; seed-wise scores were also reported.

#### Uncertainty and tests

We used 10,000 paired whole-compound bootstrap draws, holding models, folds and reference selections fixed. Each population shared one resampling matrix across registered comparisons, generated from sorted compound identifiers with NumPy seed 20261785. Two-sided tests used the centred bootstrap distribution. One-sided tests counted draws on the unfavourable side of the null. Finite-sample correction added one to numerator and denominator. More than 1% undefined draws made a formal test undetermined. Otherwise, undefined draws counted as unfavourable in tests and were omitted from percentile intervals.

Separate families compared BioMeme with the reference-informed and same-head MLPs, kernel memory with linear and MLP learners, and BioMeme with kernel memory. The nine memory comparisons summarize the memory-versus-learner contrasts of registered families K and P across the three screens; this summary was assembled after registration. Holm correction^[70]^ controlled the family-wise error rate within each family at two-sided *α* = 0.05. The sequential budget test asked whether BioMeme with 20 references retained at least 80% of each comparator’s full-panel improvement over the backbone. Comparators entered in order: linear, reference-informed MLP, then same-head MLP. Both endpoints had to pass before proceeding. The substitution test required five-reference BioMeme to improve both endpoints over all-reference memory. Budget tests used one-sided *α* = 0.025. Activity thresholds used strict solvent-DMSO wells, excluding blanks, pooled across the first four plates within target cell line, layout and time. The threshold was the 95th percentile of distances from their mean; direction error used calibration-compound target means. RNA ablation, profile identification, biological correspondence and coverage followed their registered families. Descriptive strata and additional analyses carry nominal intervals. Supplementary Note 5 defines test families.

#### Biological analyses

We compared the pooled feature-wise correlation between A549 Pilot 1 and U2OS dose responses with the correlation between Pilot 1 and Batch 2 A549. Their ratio normalized cross-cell correspondence by same-cell cross-batch reproducibility. A fixed 80-compound allow-list supplied both transitions at 48 h, with compound and feature identities aligned. We tested whether the ratio exceeded 0.30, using the replicate construction in Supplementary Note 1 and Supplementary Table 11. Descriptive variants removed specified feature groups or the leading response direction.

Transferability assessed compounds’ transfer agreement between MCF7 and U2OS. For each compound, target dose changes across four conditions were projected onto A549 changes. Disjoint A549 replicate halves supplied MCF7 and U2OS coefficients, then swapped roles. We averaged partial Spearman correlations across assignments, adjusting for A549 response strength and split-half reproducibility. The null permuted U2OS coefficients within a five-by-five grid of those covariate quintiles for 10,000 draws. Disjoint halves kept reference measurement noise separate between target-cell coefficients.

Mechanism and target labels came from the archived LINCS2020 compoundinfo_beta.txt file, matched by perturbation identifier (file hash in Supplementary Note 1). Biological neighbours were the five nearest distinct annotated compounds by provider-averaged kernel similarity. We measured the fraction sharing a mechanism-of-action or drug-target annotation with the query. Comparisons used matched RNA, RNA-key-block removal and shuffled RNA; block removal retained calibration geometry fitted with RNA. Coverage was the query’s largest kernel similarity to a target-cell reference, excluding calibration rows. Stable ranks defined coverage thirds within each direction and fold before scoring. A sequential test first assessed BioMeme’s advantage over the same-head MLP in the low-coverage third, then whether this advantage exceeded that in the high-coverage third.

#### Descriptive and additional analyses

Descriptive analyses covered identification, mechanism retrieval, activity, feature-group errors, increments from adding morphology memory, RNA and reference transfer in two orders and measurement prioritization. Tahoe cell-count and source-dimension sensitivities were registered descriptive analyses. LINCS target-mean noise was estimated as unbiased target-well variance divided by three. The measurement ledger (Fig. 5a) counted unique wells and L1000 signatures for calibration, references, query inputs and the reusable reference-cell screen. Prioritization ranked compounds by kernel variance, predicted response strength, MLP seed variance or random selection. Selected predictions were replaced by measured targets. We evaluated remaining error and active-compound recall. Kernel variance is one scalar per query in units of the kernel amplitude *σ*^2^, conditional on fitted coordinates and residual intercept; it excludes transfer-head uncertainty (Supplementary Notes 2 and 4).

Additional analyses assessed independent-well scoring, reference transfer, matched-input learners, budgets and fifth-plate screening (Supplementary Notes 9–12), shared variation, single-well combinations, wrong-compound references, cell counts and controls (Supplementary Tables 59–64; Supplementary Data 45). Runtime was assessed in a post hoc audit and replay (Supplementary Note 13). Further checks evaluated penalty selection, alternative single-well priors, complete-predictor decomposition, memory-weight deletion, scalar-reference controls and biological associations (Supplementary Note 14; Supplementary Data 46). Count-trajectory controls matched wrong reference compounds using A549 counts alone (Supplementary Data 47). Single-well mixing weights used nested eight-fold validation with each prior refitted; evaluation-optimal weights were separate descriptive bounds. Definitions were frozen before execution, after the original outcomes were known. Whole-compound bootstrap draws numbered 10,000; Supplementary Note 14 specifies each seed and any refitting within draws. Additional intervals are nominal and leave registered decisions unchanged. Supplementary Table 6 lists the scopes of independent checks. Complete results are in Supplementary Notes 1– 14, Supplementary Figs. 1–25 and Supplementary Tables 1–70.

#### Software and runtime

BioMeme and its neural and generative comparators were fitted on the CPU of a 16-GB Apple M1 computer (macOS 14.7.6). Implementation used Python 3.12.14, NumPy 2.5.3 and SciPy 1.18.1, with PyTorch 2.14.0 for neural comparators. Same-budget timing compared 12 serial BioMeme replays at 20 references with the archived neural workflows on the same CPU. Each process used one numerical thread; neural execution used four concurrent workers. Timers covered memory or neural fitting, own out-of-fold predictions and transfer fitting from prepared inputs through saving; shared calibration and key preprocessing preceded both. Neural totals included cross-validation, five final seeds and prescribed extension/sensitivity fits. All replayed predictions matched their archives exactly. Supplementary Note 13 defines timing boundaries; Supplementary Data 43 retains the records.

### Preregistration, locked scoring and verification

Five Open Science Framework (OSF) registrations specified protocols and supplements. U2OS, sci-Plex3 and Tahoe protocols preceded their recorded production fitting and formal query-target scoring. Supplementary Table 50 records three timing exceptions: a disclosed preview of four Tahoe source-dose contrast rows before registration, external registration of BBBC021 after scoring and U2OS coverage tests registered during fitting. The Tahoe plan already specified its tuned full-panel kernel before the preview; Supplementary Note 1 identifies the exposed drug, line and genes. Training-fold model selection followed the corresponding protocols. Main predictions and statistical definitions were not changed after scoring. Analyses are labelled preregistered (confirmatory or registered descriptive), additional or post hoc; development informed model design (Supplementary Note 5). Separately written implementations checked compound assignments, preprocessing, predictions, closed-form refits and statistics at the scopes in Supplementary Table 6.

Supplementary Tables 4, 5 and 50 record analysis categories, completion status and timing. Supplementary Note 5 lists deviations from the registered plans, including the title and abstract. Two registered U2OS analyses, a shared/specific error decomposition and a transfer-operator interpretation, were not executed (Supplementary Table 5).

## Use of AI-assisted tools

AI-assisted tools were used during manuscript preparation for language editing, proof-reading and quality-control support, and during code development, numerical analysis, verification and programmatic figure preparation. They did not generate or modify primary experimental data, and no generative image model was used to create or alter images. The authors directed the analyses, verified all AI-assisted output and take responsibility for the submitted work.

## Data availability

LINCS Cell Painting profiles and images are available as Cell Painting Gallery dataset cpg0004-lincs, with processed profiles in the LINCS Cell Painting repository. L1000 inputs came from the CLUE LINCS2020 Level 5 small-molecule release; A549 Pilot 1 RNA profiles came from Figshare 13181966, version 2. sci-Plex3 is deposited under GEO accession GSE139944. Tahoe-100M is available from Tahoe Bio; we used the chem-perturbridge pseudobulk-count product at revision 6c54c8eb, as specified in Supplementary Note 1. Image set BBBC021v1 is available from the Broad Bioimage Benchmark Collection. The BioMeme data and results archive is available at Zenodo (https://doi.org/10.5281/zenodo.24008759) and contains Supplementary Data 1–47, source data for all main and supplementary figures, derived result tables, archived predictions and verification records. A file manifest and data index identify the inputs and outputs supporting each analysis. Source data are provided with this paper. Supplementary Data 42 supplies reusable tasks with prepared inputs, separate evaluation targets, exact compound folds and reference selections, archived predictions and a scoring interface. Preregistration records and archived analysis plans are listed in Supplementary Note 5. The five OSF registrations are publicly accessible at https://osf.io/dmcb9, https://osf.io/kfv42, https://osf.io/8bnvc, https://osf.io/2cgz3 and https://osf.io/vmrfp.

## Code availability

The code package supplied for peer review as Supplementary Software contains the model implementation, evaluation code, preregistered protocols and independent-verification code, with a file-level source index. A minimal runnable package supplies processed U2OS example data, pinned dependencies and a command that fits the model, predicts and scores one complete transfer direction against archived results. The MIT-licensed package also provides a prepared-data memory interface and an accompanying LINCS tutorial for fitting, prediction, experiment-level decomposition, adding measurements and ranking candidates by uncertainty or predicted morphological activity. Numerical reproduction was checked on macOS (Apple silicon) and Linux (x86_64) with Python 3.12; version-specific validation records accompany the code. The manuscript source package includes figure-assembly scripts; the data archive supplies derived result tables and their numerical sources. The implementation runs on a standard CPU and requires no GPU. The MIT-licensed runnable BioMeme implementation is available at https://github.com/JIABI/BioMeme, release v4.22. This release preserves the scientific implementation of software version 4.21. The full evaluation and verification code are archived with the data in the BioMeme 4.22 Zenodo collection at https://doi.org/10.5281/zenodo.24008759.

## Acknowledgements

This work was supported by the CNPC Innovation Fund (grant no. 2024DQ02-0501), the Royal Society (grant no. IECNSFC233444), the Intelligent Manufacturing Longcheng Laboratory (grant no. CJ20254004) and the Youth Science and Technology Talent Promotion Project of Jiangsu Province (grant no. JSTJ-2025-137).

## Author contributions

J.B. contributed to all aspects of the study, including the independent-verification code. J.B. and C.Z. conceived the study. J.B., H.L., Y.W. and T.L. performed the computational experiments. J.B., X.B. and R.H. checked the data and results. J.T. and S.R.G. critically reviewed the manuscript. All authors contributed to writing, reviewing and editing the manuscript.

